# McIdas localizes at centrioles and controls centriole numbers through PLK4-dependent phosphorylation

**DOI:** 10.1101/2022.09.30.510086

**Authors:** Marina Arbi, Margarita Skamnelou, Spyridoula Bournaka, Sihem Zitouni, Stavroula Tsaridou, Ozge Karayel, Catherine G. Vasilopoulou, Aikaterini C. Tsika, Nikolaos N. Giakoumakis, Georgios A. Spyroulias, Matthias Mann, Mónica Bettencourt-Dias, Stavros Taraviras, Zoi Lygerou

## Abstract

The centriole duplication cycle must be tightly controlled and coordinated with the chromosome cycle. Aberrations in centriole biogenesis can lead to cancer, developmental disorders and ciliopathies. Here, we show that McIdas -previously implicated in cell cycle control and centriole amplification in multiciliated cells-is critical to maintain centriole numbers. Using expansion microscopy, we demonstrate that McIdas is present at the middle part of centrioles, where it exhibits a differential localization during the cell cycle. McIdas loss perturbs daughter centriole biogenesis and centrosomal SAS6 recruitment, whereas its overexpression induces centriole overduplication. Consistently, McIdas depletion reduces PLK4-induced centriole amplification. McIdas interacts with and is phosphorylated by PLK4 in multiple sites identified by mass spectrometry. Mutational analysis shows that McIdas phosphorylation is important for centriole number control. Overall, our results identify a novel, direct role of McIdas on centriole duplication that can link its previously characterized roles in the chromosome cycle and multiciliogenesis.

## Introduction

Centrioles are highly conserved, microtubule-based cylindrical organelles that contribute to the biogenesis of centrosomes and primary or motile cilia in cells (Jana, 2021; Paz and Luders, 2018; Vasquez-Limeta and Loncarek, 2021; Wu and Akhmanova, 2017). Two orthogonally arranged centrioles, one mother and one daughter, surrounded by the pericentriolar material (PCM), are the major components of centrosomes. Centrosomes play critical roles in cell division, cell migration, polarization and signaling. In many non-dividing cells, the centrosome migrates to the apical cell membrane, where the mature mother centriole docks and acting as basal body nucleates a primary, sensory immotile cilium (Arslanhan et al., 2020). However, in terminally differentiated multiciliated cells centrioles are massively amplified and act as templates for the generation of multiple motile cilia (Meunier and Azimzadeh, 2016; Spassky and Meunier, 2017).

Centrosomes duplicate once and only once during the cell cycle, in S phase. Licensing, duplication and segregation are key steps that control the centriole duplication cycle (Firat-Karalar and Stearns, 2014; Fu et al., 2015; Gomes Pereira et al., 2021). Upon exit from M phase and during G1 phase, the two centrioles are disengaged, licensing a new round of centriole duplication. During S phase, one new centriole -called procentriole-is orthogonally generated next to each pre-existing centriole. The newly generated centrioles are elongated during G2 phase, while mother centrioles mature. During mitosis, the two pairs of centrioles are separated to form the poles of the mitotic spindle, ensuring accurate sister chromatid segregation to the two daughter cells. PLK4 (Polo-like kinase 4), STIL (SCL/TAL1 interrupting locus protein) and SAS6 (Spindle assembly abnormal protein 6 homolog) are critical regulators for the initiation of centriole assembly (Arquint and Nigg, 2016). PLK4 is first recruited as a ring-like structure at the proximal region of mother centrioles (Bettencourt-Dias et al., 2005; Habedanck et al., 2005; Kim et al., 2013; Ohta et al., 2014). PLK4 binds and phosphorylates STIL, resulting in the subsequent loading of SAS6 (Kratz et al., 2015; Moyer et al., 2015; Ohta *et al*., 2014). This leads to the assembly of the cartwheel, a macromolecular hub nucleating the formation of the ninefold symmetrical microtubule triplets of the new centriole (Gonczy, 2012). The expression levels and the interactions between PLK4, STIL and SAS6 must be strictly regulated with their overexpression leading to the formation of multiple daughter centrioles around a mother centriole in rosette-type structures. PLK4 is regulated through multiple post-transcriptional and post-translational mechanisms (Cunha-Ferreira et al., 2013; Cunha-Ferreira et al., 2009; Guderian et al., 2010; Holland et al., 2012; Holland et al., 2010; Lopes et al., 2015; Nakamura et al., 2013; Phan et al., 2022; Rogers et al., 2009; Ryniawec and Rogers, 2022), whereas STIL and SAS6 are degraded in late M and early G1 phase by the anaphase-promoting complex/cyclosome associated with Cdh1 (APC/C^Cdh1^) (Arquint and Nigg, 2014; Arquint et al., 2012; Strnad et al., 2007).

The centriole cycle must be closely coordinated with the chromosome cycle, as defects in this coordination will lead to improper chromosome segregation, causing genomic instability and tumorigenesis (Gonczy, 2015; Nigg and Holland, 2018; Raff and Basto, 2017). Earlier studies showed that cyclin-dependent kinases are required for centrosome duplication (Hinchcliffe et al., 1999; Lacey et al., 1999; Matsumoto et al., 1999; Meraldi et al., 1999; Zitouni et al., 2016). Moreover, replication proteins such as MCM5, Orc1, Geminin and Cdc6 are recruited at centrosomes through interactions with S phase cyclins and are implicated in centriole duplication (Ferguson and Maller, 2008; Ferguson et al., 2010; Hemerly et al., 2009; Tachibana et al., 2005; Xu et al., 2017). However, how the centriole and chromosome cycles are inter-linked remains poorly characterized.

Recent studies proposed McIdas as a protein with roles both in cell cycle control and in differentiation decisions in cells (Arbi et al., 2018). McIdas is a nuclear, vertebrate-specific coiled-coil protein (Pefani et al., 2011), phylogenetically related to the cell cycle regulators Geminin (McGarry and Kirschner, 1998) and GemC1 (Balestrini et al., 2010). McIdas binds to Geminin and inhibits its association with Cdt1, affecting DNA replication licensing (Pefani *et al*., 2011). Its protein levels drop during anaphase of mitosis, and it has been suggested as an APC/C target. McIdas loss affects normal cell cycle progression and leads to the accumulation of cells in S phase, whereas its overexpression causes abnormal multipolar spindles (Pefani *et al*., 2011). In addition, several studies identified both McIdas and GemC1 as early regulators of multiciliogenesis (Arbi et al., 2016; Boon et al., 2014; Kyrousi et al., 2015; Stubbs et al., 2012; Terre et al., 2016; Zhou et al., 2015). During multiciliogenesis, McIdas acts through the formation of complexes with transcription factors, including E2F4/5 and DP1, switching on a gene expression program essential for centriole amplification and cilia formation (Ma et al., 2014).

Here, we show that McIdas is a novel regulator of the centriole duplication cycle. McIdas localizes at centrioles in a cell cycle specific manner, interacts with and is phosphorylated by PLK4 and is required for the maintenance of correct centriole numbers. Our data suggest that the roles of McIdas in the chromosome cycle and multiciliogenesis may be interconnected, through a novel, direct role of McIdas in centriole duplication.

## Results

### McIdas is localized at centrosomes in a cell cycle specific manner

McIdas was initially characterized as a nuclear protein which binds to Geminin and inhibits its association with Cdt1 (Pefani *et al*., 2011). Overexpression of McIdas causes abnormal mitotic spindle formation (multipolar spindles), indicating that it may affect centrosome numbers in cycling cells. Moreover, McIdas was identified as an important regulator of centriole and cilia formation in post-mitotic multiciliated cells (Stubbs *et al*., 2012). It exhibits a highly specific expression pattern in ciliated epithelia, but it is also expressed in proliferating cells, albeit at lower levels (Arbi *et al*., 2016). To gain insight into a possible localization of McIdas at centrosomes, we followed its endogenous expression in human osteosarcoma U2OS cells, using a specific antibody against human McIdas protein (Pefani *et al*., 2011). Both EdU incorporation and DNA staining were used to monitor the different cell cycle phases, whereas an antibody against Centrin was used to mark centrioles. As shown in Figure 1A, during G1 phase of the cell cycle one McIdas dot can be detected at centrosome adjacent to two Centrin dots, whereas from early S phase McIdas was recruited at both parental centrioles, as two McIdas dots were apparent. Its expression at centrosomes increases as cell cycle progresses and drops at the end of mitosis, during telophase. This is consistent with a previous study showing that McIdas total protein levels drop during anaphase (Pefani *et al*., 2011). A GFP-tagged McIdas protein was also detected at centrosomes in different cell lines, either as an exogenously overexpressed protein (Supplementary Figure 1A & B) or stably expressed in HeLa cells (Figure 1B). Accumulation of Centrin was evident upon GFP-McIdas overexpression in U2OS and hTERT-RPE1 cells, suggestive of a possible role of McIdas in centriole formation (Supplementary Figure 1A & B, marked by an asterisk). Notably, addition of the proteasome inhibitor MG132 in HeLa cells stably expressing either GFP-tagged McIdas or GFP alone as a control, blocked McIdas degradation, causing its accumulation at centrosomes (Supplementary Figure 1C). The specificity of McIdas localization at centrosomes was confirmed by using two different siRNA oligos (Supplementary Figure 1D-G). Both siRNA oligos efficiently reduced McIdas expression at centrosomes and thus were used interchangeably in the following experiments. All the above data show that McIdas is expressed at centrosomes and its expression is regulated during the cell cycle. Interestingly, McIdas temporal expression pattern is reminiscent of the expression pattern followed by known centriole regulators, such as STIL and SAS6 (Arquint and Nigg, 2014; Arquint *et al*., 2012; Strnad *et al*., 2007).

**Figure 1.**
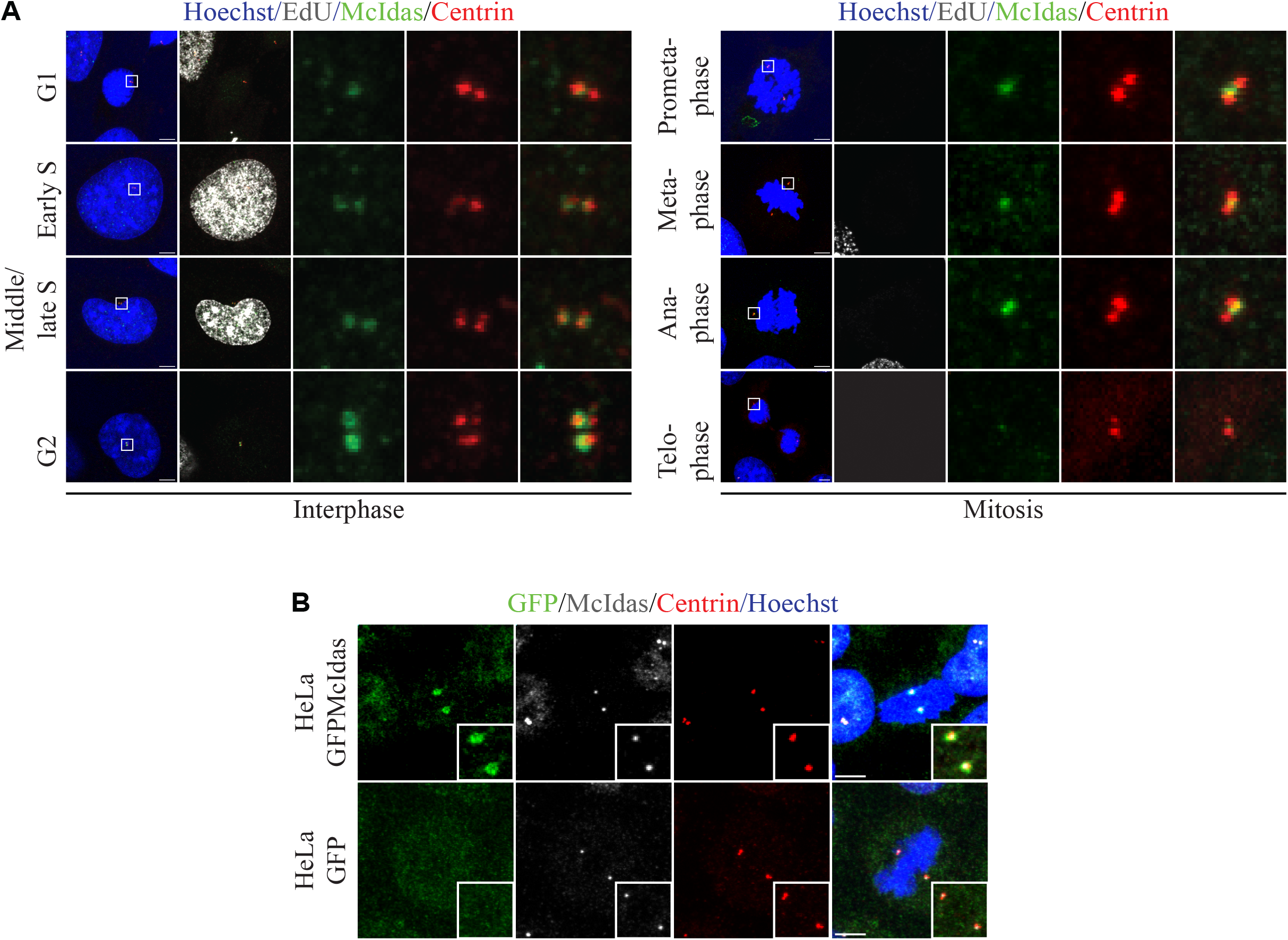
McIdas localizes at centrosomes in a cell cycle specific manner. (A) U2OS cells were immunostained with antibodies against McIdas and Centrin (distal lumen centriole marker). EdU (marks S phase cells) and DNA stains (Hoechst) were used to monitor the different cell cycle stages. Inserts correspond to the higher magnification images shown on the right for McIdas and Centrin stains at centrosomes. Arrow indicates the initial McIdas signal at centrosome during G1 phase. (B) HeLa cells were generated to stably express either GFP-tagged McIdas or GFP alone as a control. Cells were pre-extracted and immunostained with antibodies against GFP, to mark the transfected cells, endogenous McIdas and Centrin. Inserts indicate higher-magnification images of the indicated stains at centrosomes. DNA was stained with Hoechst. Scale bars, 5 μm.

### McIdas localizes at the middle of the centriole and shows a differential localization pattern during the cell cycle

Given that McIdas was initially detected at centrosome as only one dot during G1 phase, we next examined in which one of the two centrioles it can be found. To that end, we performed an immunofluorescence analysis in hTERT-RPE1 cells (also in U2OS cells, data not shown), stained for McIdas along with Cep164, which marks the distal appendages of mother centriole. As shown in Figure 2A, McIdas was initially expressed as one dot in the Cep164-positive centriole. To study McIdas sub-centrosomal localization, further analysis was performed by comparing McIdas expression with additional centriole markers, such as Centrin (distal centriole lumen marker) and Cep135 (proximal marker). McIdas-, Cep135- and Cep164-signal intensities were plotted relative to distance and revealed that McIdas localization at centrosome was proximal to that of Centrin and Cep164 and distal to Cep135 (Figure 2B & C). McIdas is partially co-localized with Cep164 at the distal part of centrioles. Notably, following McIdas expression at the newly synthesized centrioles during early mitosis, we observed that McIdas expression is significantly increased at procentrioles as cell cycle progresses, while it becomes significantly reduced at mother centrioles (Figure 2D). To corroborate this result, we followed McIdas localization in respect to SAS6, a component of cartwheel structure in the proximal lumen of daughter centrioles. We found that indeed during G2 phase McIdas is mainly localized at the newly synthesized centrioles, as McIdas can be found at the distal part of SAS6 positive centriole, whereas it cannot be detected anymore at the distal part of mother centriole (Figure 2E-I). Interestingly, McIdas expression at centrioles preceded the expression of SAS6, highlighting an early recruitment of McIdas at centrioles during centriole duplication.

**Figure 2.**
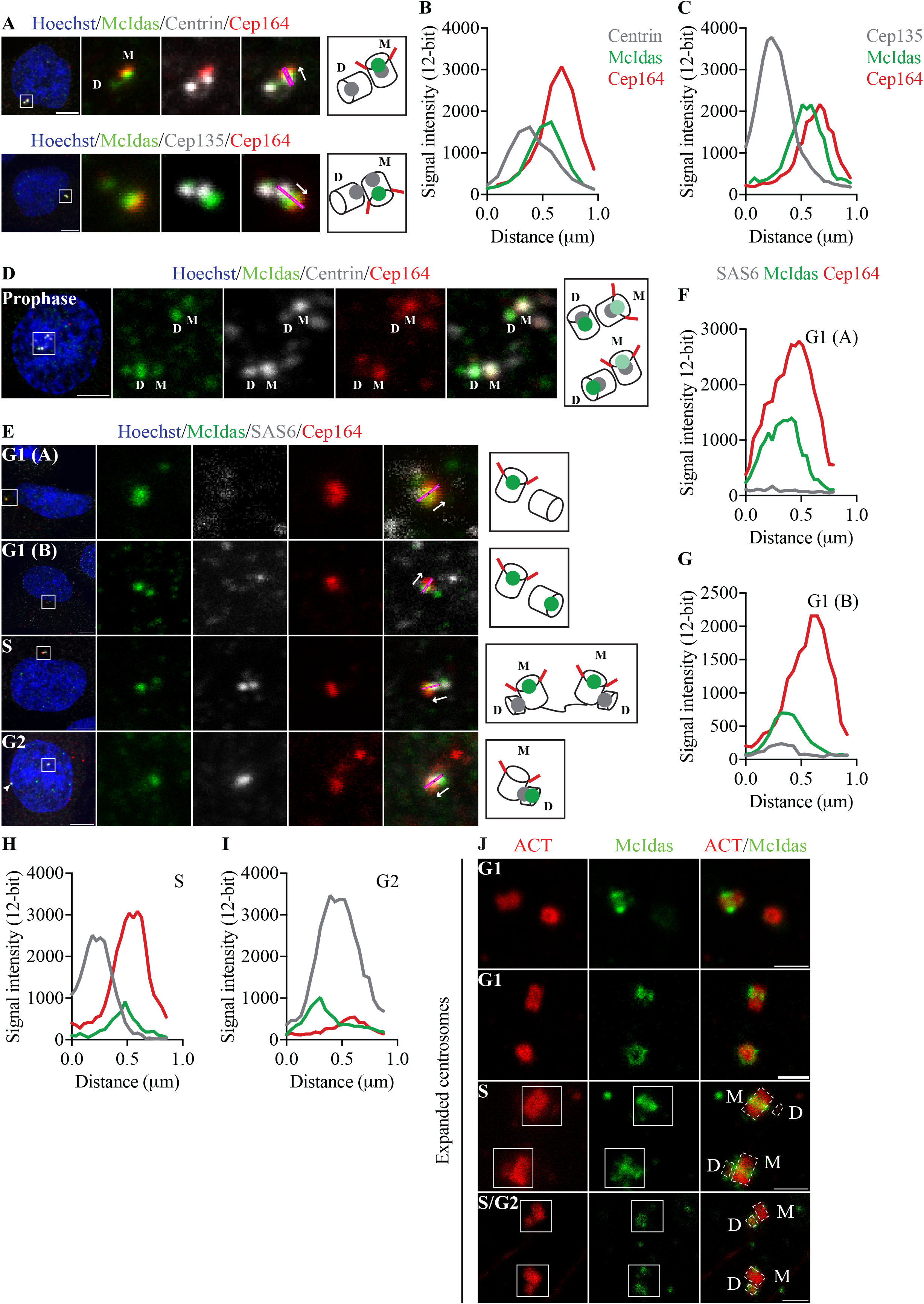
McIdas localizes at the middle part of centrioles and relocalizes from mother to daughter centriole during G2 phase. (A) hTERT-RPE1 cells were stained for McIdas and different centriole markers, including Cep135 (proximal centriole marker) or Centrin (distal lumen centriole marker) and Cep164 (distal appendage centriole marker). Inserts correspond to the higher-magnification images shown on the right for the indicated stains at centrosomes. (B, C) Corresponding fluorescent intensity plots of the centrosomal signals of McIdas, Cep164 and Centrin or Cep135, respectively. Position and direction of the line used to generate the intensity profiles as a function of distance are indicated at (A) by the magenta line and the arrow, respectively. (D) Representative image from a prophase hTERT-RPE1 cell showing duplicated centrosomes that have been stained with antibodies against McIdas, Centrin and Cep164. Insert corresponds to the higher-magnification images of the indicated stains at centrosomes shown on the right. (E) hTERT-RPE1 cells at different cell cycle stages were immunostained with antibodies against McIdas, SAS6 (proximal lumen of daughter centrioles) and Cep164. SAS6 and Cep164 expression patterns were used to classify the cells at the different cell cycle phases. Arrowhead corresponds to the second centrosome of this G2-phase cell. (F-I) Corresponding fluorescent intensity plots of the centrosomal signals of McIdas, SAS6 and Cep164 analyzed at (E). Position and direction of the line used to generate the intensity profiles are indicated at (E) by the magenta line and the arrow, respectively. (J) ExM images from hTERT-RPE1 cells at different cell cycle phases stained with antibodies against acetylated α-tubulin (ACT, centriolar MT marker) and McIdas. Centriole numbers and acetylated α-tubulin staining were used to identify the different cell cycle phases. Mother and newly synthesized centrioles are outlined with dashed lines. Schematic representations at A, D & E indicate McIdas localization with respect to centriole markers used, that would be consistent with this analysis. DNA was stained with Hoechst. Scale bars, 5 μm (A, D & E) or 2 μm (J). ExM ∼ 4x Abbreviations: ACT: acetylated α-tubulin, M: mother centriole, D: daughter centriole, AU: arbitrary units.

To gain a better insight into this sub-centrosomal localization of McIdas we performed expansion microscopy (ExM), marking centrioles with acetylated α-tubulin (Figure 2J). In line with our previous analysis, McIdas is localized at the middle part of mother centrioles and exhibits one- or two-dots staining in G1 phase-cells, and this localization was consistent with McIdas distribution on the outer surface of centrioles. During late S and G2 phases of the cell cycle, higher resolution images verified the gradual enrichment of McIdas on the outer surface of the newly synthesized centrioles, as opposed to decreased levels on mother centrioles.

We therefore conclude that McIdas is localized at the middle part of centrioles, displaying enrichment from mother to daughter centrioles as cell cycle progresses.

### McIdas regulates daughter centriole biogenesis

Given that McIdas localizes at centrosomes we next manipulated its expression in cells to investigate McIdas role on centriole biogenesis. McIdas was initially characterized as a cell cycle regulator (Pefani *et al*., 2011). To study its direct role on centrosome cycle, we performed a centrosome duplication assay in U2OS cells where centrosome cycle can be experimentally uncoupled from the DNA replication cycle by inhibiting DNA synthesis (Meraldi *et al*., 1999). To assess this, U2OS cells were treated with high concentration of hydroxyurea (HU), resulting in multiple rounds of centriole duplication during a prolonged S-phase. First, to examine whether McIdas is sufficient to induce centriole amplification we treated the cells with HU and overexpressed either a GFP-McIdas fusion protein or GFP alone as a control. FACS analysis did not detect significant differences in the cell cycle profile of these cells, suggesting that the differences in centriole numbers is cell-cycle independent (data not shown). We observed that 34% of the control GFP-overexpressing cells possess more than 4 centrioles (CP110 dots per cell) due to HU treatment, whereas 60% of the GFP-McIdas overexpressing cells possess more than 4 centrioles (Figure 3A and B). The above results highlight that increased McIdas expression is sufficient to directly induce centriole amplification in arrested cells.

**Figure 3.**
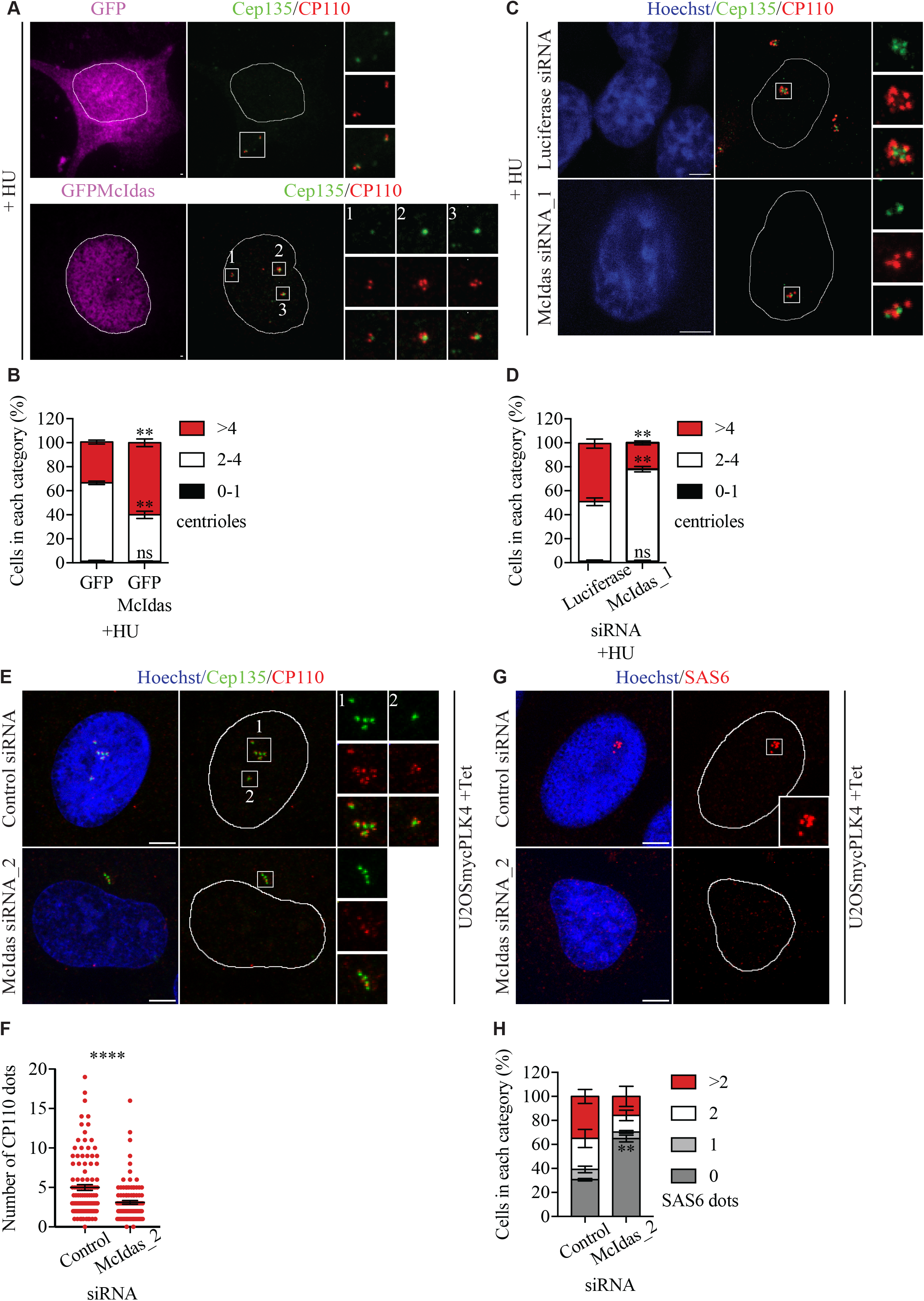
McIdas is essential for centriole amplification. (A) U2OS cells were treated with 4mM hydroxyurea (HU) and 24 h later were transfected with vectors expressing GFP-tagged McIdas or GFP alone as a control. Cells were fixed with methanol 48 h after the transfections and centriole numbers were counted by using antibodies against Cep135 and CP110, to mark the proximal and distal centriole parts, respectively. Inserts correspond to the higher-magnification images of the indicated stains at centrosomes shown on the right. (B) Quantification of centriole numbers per cell in GFP- and GFP-McIdas-overexpressing cells. Data are presented as the mean values of three independent experiments and error bars indicate ± SEM. In each experiment at least 100 cells were counted per condition. (C) McIdas siRNA_1 or control siRNA oligos were transfected in U2OS cells. 24 h later, after a second transfection with the indicated siRNAs, cells were treated with 4mM HU for another 48 h. Cells were fixed and centriole numbers were determined by staining with antibodies against Cep135 and CP110. Inserts correspond to the higher-magnification images shown on the right of Cep135 and CP110 stains at centrosomes. (D) Quantification of centriole numbers per cell in control and McIdas-depleted cells. Data are presented as the mean values of three independent experiments and error bars indicate ± SEM. In each experiment at least 100 cells were counted per condition. (E, G) U2OS cells expressing myc-PLK4 under a tetracycline-dependent promoter were transfected with McIdas siRNA_2 or control siRNA oligos. A second transfection with the indicated siRNAs followed 24 h later and then cells treated with tetracycline for an efficient myc-PLK4 overexpression. Cells were fixed and immunostained for Cep135 and CP110 to count centriole numbers (E) or SAS6 (G). Inserts correspond to higher-magnification images of the indicated stains at centrosomes. (F) Representative quantification of centriole numbers in control and McIdas-depleted cells, overexpressing PLK4. At least two independent experiments have been performed. In each experiment more than 100 cells were counted per condition. (H) Quantification of SAS6 foci in control and McIdas-depleted cells, overexpressing PLK4. Data are presented as the mean values of two independent experiments and error bars indicate ± SEM. In each experiment at least 100 cells were counted per condition. Nuclei boundaries are marked with lines. DNA was stained with Hoechst. Scale bars, 5 μm. *P*-values in B, D and H were calculated using two-tailed Student’s t-test. *P*-value in F was calculated by the nonparametric two-tailed Mann-Whitney test. \*\**P* < 0.01, \*\*\*\**P* < 0.0001 Abbreviations: HU: hydroxyurea, tet: tetracycline, ns: not significant.

Next, we depleted the endogenous protein from U2OS cells, followed by treatment with hydroxyurea (HU) and we determined the number of centrioles by staining the cells with markers for the proximal and distal ends of centrioles (Cep135 and CP110, respectively). McIdas was efficiently depleted from U2OS cells as assessed by real-time PCR experiments (Supplementary Figure 2A). As shown in Figure 3C and D, almost half of the control cell population possessed more than 4 centrioles due to HU treatment, whereas after McIdas silencing only 22% of cells exhibited more than 4 centrioles. By this analysis we found that centrosome overduplication caused by HU was impaired in McIdas-depleted cells. Given that PLK4 overexpression induces centriole amplification we next examined whether McIdas can also affect PLK4-mediated centriole amplification. To that end, we depleted McIdas in U2OS cells inducibly overexpressing PLK4 and we measured the number of centrioles per cell, by using either CP110 or Centrin as centriole markers. McIdas was efficiently depleted from U2OS cells as assessed by immunoblotting and real-time PCR experiments (Supplementary Figure 2B & C). PLK4-overexpressing cells possess an increased number of cells with amplified centrioles organized in rosette-like structures, whereas this phenotype was severely reduced in McIdas-knockdown cells as revealed by CP110 and Centrin staining (Figure 3E & F, Supplementary Figure 2D & E). Moreover, we examined the effect of McIdas depletion on cartwheel assembly in PLK4-overexpressing cells. In control cells there is an increased recruitment of McIdas along with PLK4 and SAS6 in the rosette-like structures (Supplementary Figure 2F & G). On the contrary, there is a significant increase of cells without SAS6 foci upon McIdas depletion, highlighting that McIdas is important for centrosomal SAS6 recruitment (Figure 3G & H).

We further tested whether McIdas is also required for canonical centriole duplication, by depleting its expression in asynchronous U2OS cells (Figure 4). McIdas was efficiently depleted from U2OS cells as assessed by real-time PCR experiments (Supplementary Figure 2H). We focused on S phase cells (EdU positive cells), and we counted the number of Centrin foci, which corresponds to the number of centrioles. Most of the control cells possess 4 centrioles as centriole duplication takes place during S phase, whereas upon McIdas depletion there is a statistically significant increase in the number of cells with 2 Centrin dots, highlighting unduplicated centrosomes (Figure 4A & B). Moreover, we investigated whether McIdas functions during cartwheel assembly, following SAS6 recruitment at centrosomes. To that end, we depleted McIdas from U2OS (and RPE1 cells, data not shown) asynchronous cells and counted the number of SAS6 foci in EdU positive cells, where 2 SAS6 foci were expected to be found at centrosomes. Upon McIdas depletion there was a significant increase in cells with 0 or 1 SAS-6 focus, as compared to control EdU positive cells where most cells possess two SAS6 foci (Figure 4C & D). This result indicates that SAS6 recruitment at centrosomes is perturbed upon McIdas loss.

**Figure 4.**
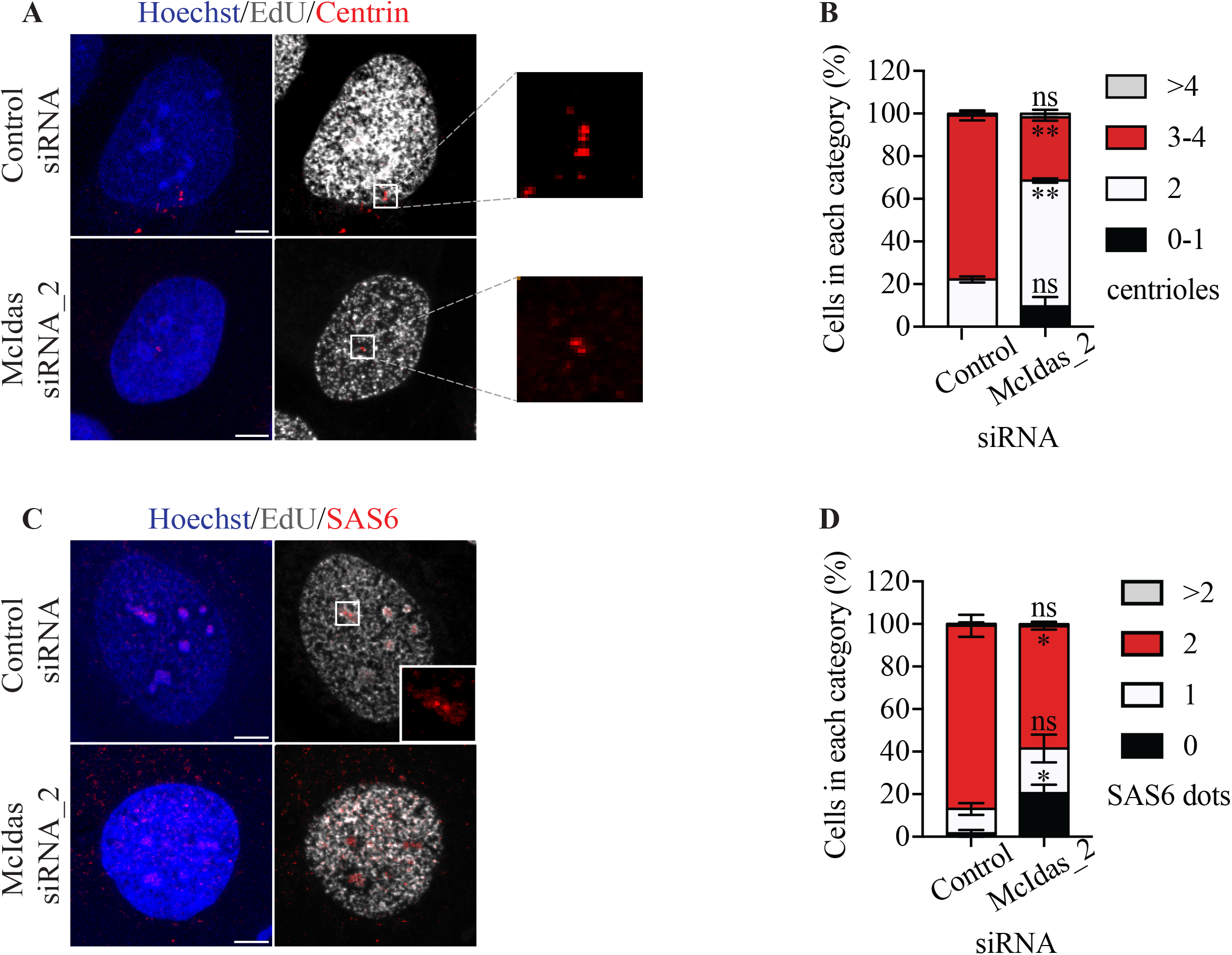
McIdas is required for daughter centriole biogenesis in cycling cells. (A, C) U2OS cells were transfected with McIdas siRNA_2 or control siRNA oligos and were immunostained with antibodies against Centrin to count centriole numbers (A) or SAS6 (C). Inserts correspond to higher-magnification images of the indicated stains at centrosomes. (B) Quantification of centriole numbers per cell in control and McIdas-depleted cells. Data are presented as the mean values of two independent experiments and error bars indicate ± SEM. In each experiment at least 100 cells were counted per condition. (D) Quantification of SAS6 foci in control and McIdas-depleted cells. Data are presented as the mean values of two independent experiments and error bars indicate ± SEM. In each experiment at least 100 cells were counted per condition. DNA was stained with Hoechst. Scale bars, 5 μm. *P*-values were calculated using two-tailed Student’s t-test. \**P* < 0.1, \*\**P* < 0.01. Abbreviations: ns: not significant.

Altogether, the above data strongly support that McIdas expression is important for the maintenance of correct centriole numbers in cancer and normal cycling and S-phase arrested cells, affecting SAS6 recruitment, an essential component for cartwheel assembly.

### McIdas interacts with and is phosphorylated by PLK4

To investigate how McIdas acts molecularly during centriole duplication we first asked whether McIdas can interact with the core centriole duplication machinery, including interactions with PLK4, STIL and SAS6 proteins. To that end, we immunoprecipitated GFP-McIdas or GFP alone as a control from U2OS cells expressing myc-PLK4 under a tetracycline-dependent promoter and we found that PLK4 was present in McIdas immunoprecipitates (Figure 5A). Interestingly, we were not able to detect interactions between McIdas and STIL (Supplementary Figure 3A & B) or SAS6 proteins (Supplementary Figure 3C & D).

**Figure 5.**
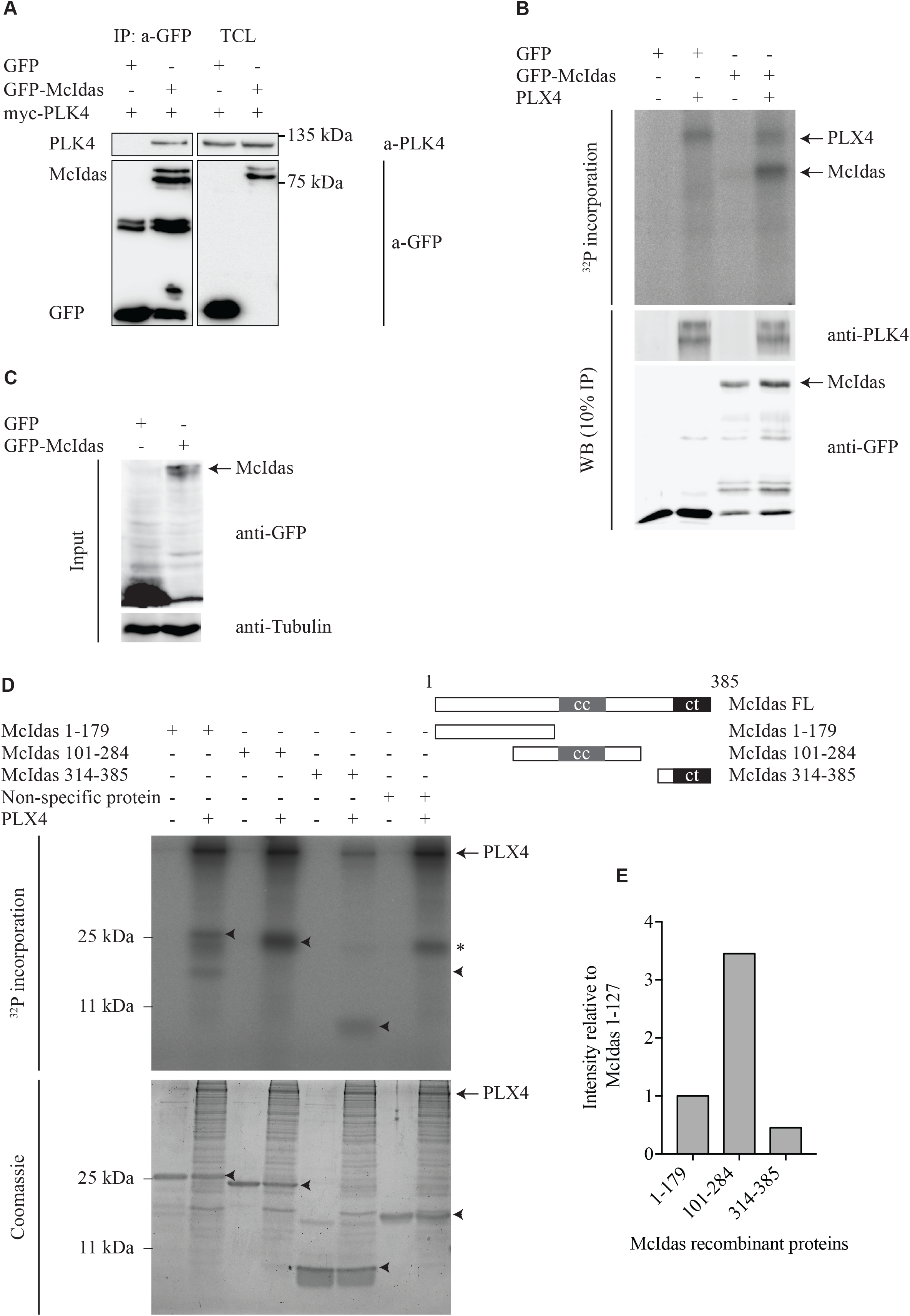
McIdas interacts with and is phosphorylated by PLK4 at multiple sites. (A) GFP-tagged McIdas or GFP alone as a control were transfected in an inducible U2OS myc-PLK4 overexpressing cell line. Immunoprecipitation was performed against GFP and PLK4 was detected in McIdas immunoprecipitates. (B) *In vitro* kinase assays were performed by using IP control or GFP-McIdas, incubated with recombinant Xenopus PLK4 (PLX4) and [γ-^32^P] ATP. Samples were analyzed by autoradiography. Western blot analysis was performed in the 10% of each sample used in kinase assays, by staining with antibodies against PLK4 and GFP to follow loaded proteins. (C) Western blot analysis from total cell extracts used for the *in vitro* kinase assays performed at (B) by using antibodies against GFP and tubulin. (D) *In vitro* kinase assays were performed by using three truncated McIdas recombinant proteins as indicated. Each reaction was incubated with recombinant Xenopus PLK4 (PLX4) and [γ-^32^P] ATP and samples were analyzed by autoradiography. Coomassie staining was used to follow the loaded proteins. A non-specific protein was used as a negative control. Arrowheads indicate the expected position of the proteins. Asterisk represents a non-specific band. (E) Representative quantification of phosphorylation of McIdas recombinant proteins used at (D), calculated by Image Lab software. Values indicate phosphorylation signal intensity relative to McIdas 1-179 recombinant protein (set as 1). At least three independent experiments have been performed.

Given the specific interaction between McIdas and the kinase PLK4 we further examined whether McIdas can be a substrate of PLK4. First, *in vitro* kinase assays were performed by incubating the immunoprecipitated GFP-McIdas from HEK293T cells with recombinant PLK4 and ^32^P-radiolabeled ATP. Our analysis showed incorporation of ^32^P-radiolabeled ATP into McIdas in the presence of PLK4 (Figure 5B). Thus, McIdas interacts with PLK4 and is a novel substrate of PLK4.

Moreover, to determine which part of McIdas protein mediates its phosphorylation by PLK4 we performed *in vitro* kinase assays by using three truncated McIdas recombinant proteins, including an N-terminal fragment of McIdas (1-179 aa), a fragment containing the central coiled-coil domain (101-284 aa) and a fragment containing the conserved C-terminal region of McIdas (314-385 aa). Each one of these proteins was incubated with purified PLK4 in the presence of ^32^P-radiolabeled ATP. Consistently, we observed that McIdas is indeed phosphorylated by PLK4, but in multiple sites, with an increased incorporation of ^32^P-radiolabeled ATP on its coiled-coil containing fragment (Figure 5D & E). An irrelevant recombinant protein which couldn’t be phosphorylated was used here as a negative control. All the above results suggest that PLK4 binds and phosphorylates McIdas at multiple sites.

Next, we performed a tandem mass spectrometry analysis to map the PLK4-specific phosphorylation sites on McIdas protein. To that end, *in vitro* phosphorylated McIdas fragments were subjected to mass spectrometry and 10 serine phosphorylation sites were identified (Figure 6A). Among those sites, 4 of them (S55, S58 and S86/87) are located to the N-terminal region of McIdas, 3 are located on both sides of the central coiled-coil region (S167, S246, S271), whereas the last 3 are found in the C-terminal region of the protein (S333, S338, S340) (Figure 6B). Next, we mutated these 10 serine residues into alanine (GFP-McIdas 10A) or aspartic acid (GFP-McIdas 10D) residues to generate phospho-dead or phospho-mimetic mutant forms of McIdas protein, respectively. We first overexpressed either GFP-McIdas WT or GFP-McIdas 10A in HEK293T cells. These proteins were immunoprecipitated from cells and incubated with purified PLK4 and ^32^P-radiolabeled ATP for an *in vitro* kinase assay. The results revealed that the phospho-dead mutant of McIdas exhibits a severely impaired incorporation of ^32^P-radiolabeled ATP, suggesting the importance of the identified sites for McIdas phosphorylation (Figure 6C & E). Interestingly, as shown in Figure 6D and Supplementary Figure 4D, the slower mobility shift band of McIdas WT suggests a phosphorylated form compared to the faster mobility shift band of the phospho-dead mutant that represents a non-phosphorylated McIdas form. Consistently, incubation of GFP-McIdas WT- or 10A- or 10D-containing extracts with lambda protein phosphatase released these proteins from their phosphate groups, suggesting phosphorylation as the responsible modification for the observed differences on their mobility (Supplementary Figure 4A). To narrow down the serine residues that could be important for McIdas phosphorylation by PLK4, we generated partial phospho-dead McIdas mutants (GFP-McIdas 4A_N or 3A_cc or 3A_C mutant forms) (Supplementary Figure 4B). The above analysis shows that multiple serine residues were phosphorylated by PLK4, with a greater preference in the serine residues that are located on both sides of the coiled-coil region (Supplementary Figure 4C & E). This is consistent with our previous analysis showing increased phosphorylation of McIdas on its coiled coil-containing region (Figure 5D & E).

**Figure 6.**
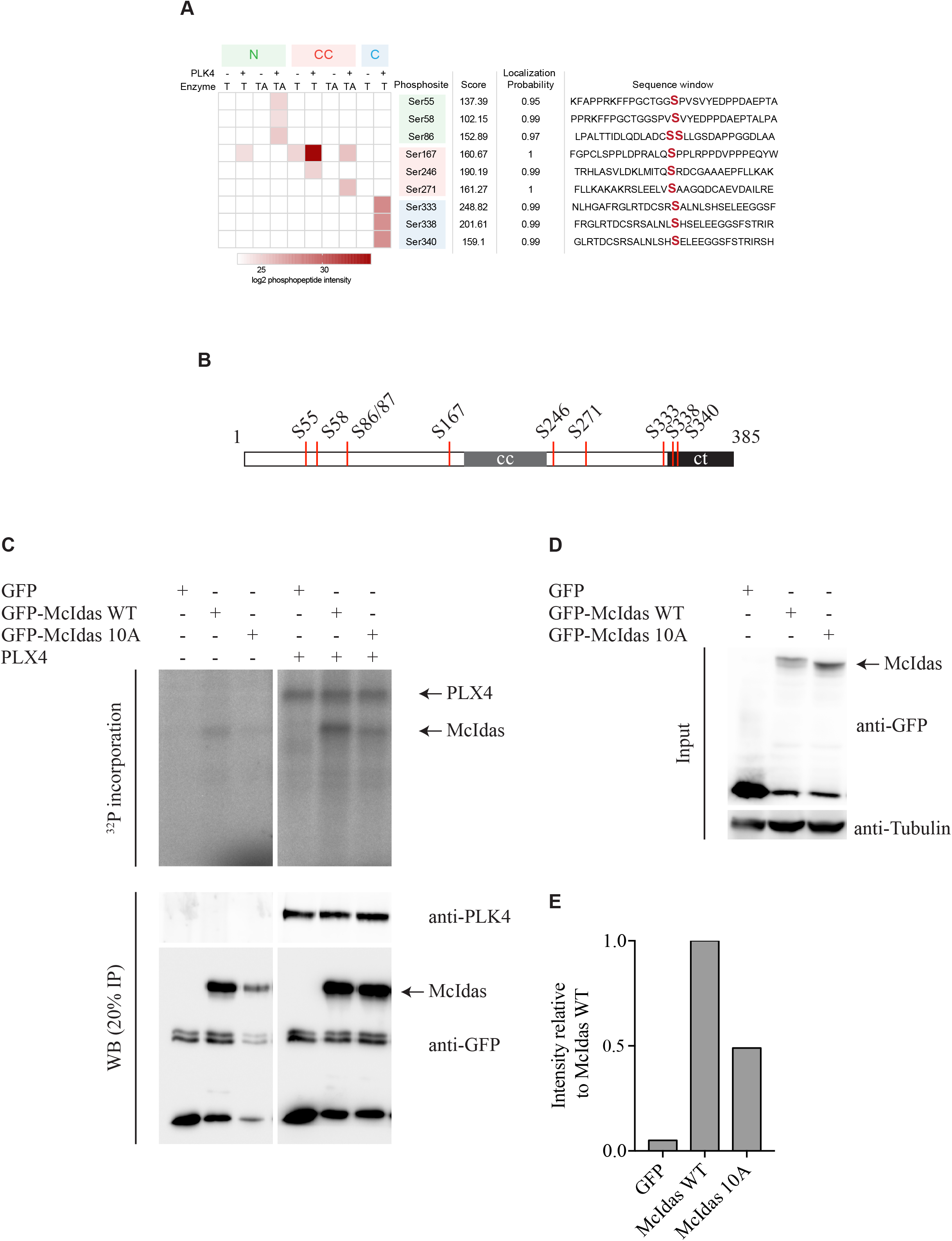
PLK4 phosphorylates McIdas at 10 serine residues. (A) *In vitro* phosphorylated McIdas recombinant proteins were subjected to tandem mass spectrometry analysis and 10 PLK4-specific serine phosphorylations were identified. (B) Schematic representation of the PLK4-phosphorylated serine residues identified on McIdas protein. (C) *In vitro* kinase assays were performed using IP control or GFP-McIdas WT or the phospho-dead GFP-McIdas 10A. Each IP was incubated with recombinant Xenopus PLK4 (PLX4) and [γ-^32^P] ATP. Samples were analyzed by autoradiography. Western blot analysis was performed in the 20% of each sample by using antibodies against PLK4 and GFP to monitor loaded proteins. (D) Western blot analysis of total cell extracts used at (C) for the *in vitro* kinase assays was performed by using antibodies against GFP and tubulin. (E) Representative quantification of phosphorylation of McIdas WT or 10A mutant used at (C), calculated by Image Lab software. Values indicate phosphorylation signal intensity relative to McIdas WT (set as 1). At least three independent experiments have been performed.

We therefore conclude that McIdas interacts with PLK4 and is a novel substrate of PLK4. Our data identified multiple PLK4-specific phosphorylation sites on McIdas protein that could be important for McIdas function at centrosomes.

### PLK4-specific phosphorylation of McIdas is essential for daughter centriole biogenesis

To examine the significance of McIdas phosphorylation by PLK4 on centriole duplication, U2OS cells were treated with HU and transfected with plasmids expressing either GFP-McIdas WT gene or 10A and 10D mutants. Following quantification of centrioles by using CP110 marker, we observed that the GFP-McIdas 10A phospho-dead mutant cannot induce centriole amplification upon HU treatment, as compared to GFP-McIdas WT (Figure 7A and B). This analysis points out that McIdas phosphorylation by PLK4 is a regulatory mechanism essential for centriole biogenesis. Interestingly, the phospho-mimetic mutant GFP-McIdas 10D, which represents a continuously phosphorylated protein, was also not able to induce centriole amplification as compared to GFP-McIdas WT-overexpressing cells. This result suggests that constant phosphorylation of McIdas by PLK4 also inhibits centriole biogenesis. We therefore conclude that both McIdas phosphorylation and dephosphorylation could be important mechanisms that act antagonistically for centriole number control in cells.

**Figure 7.**
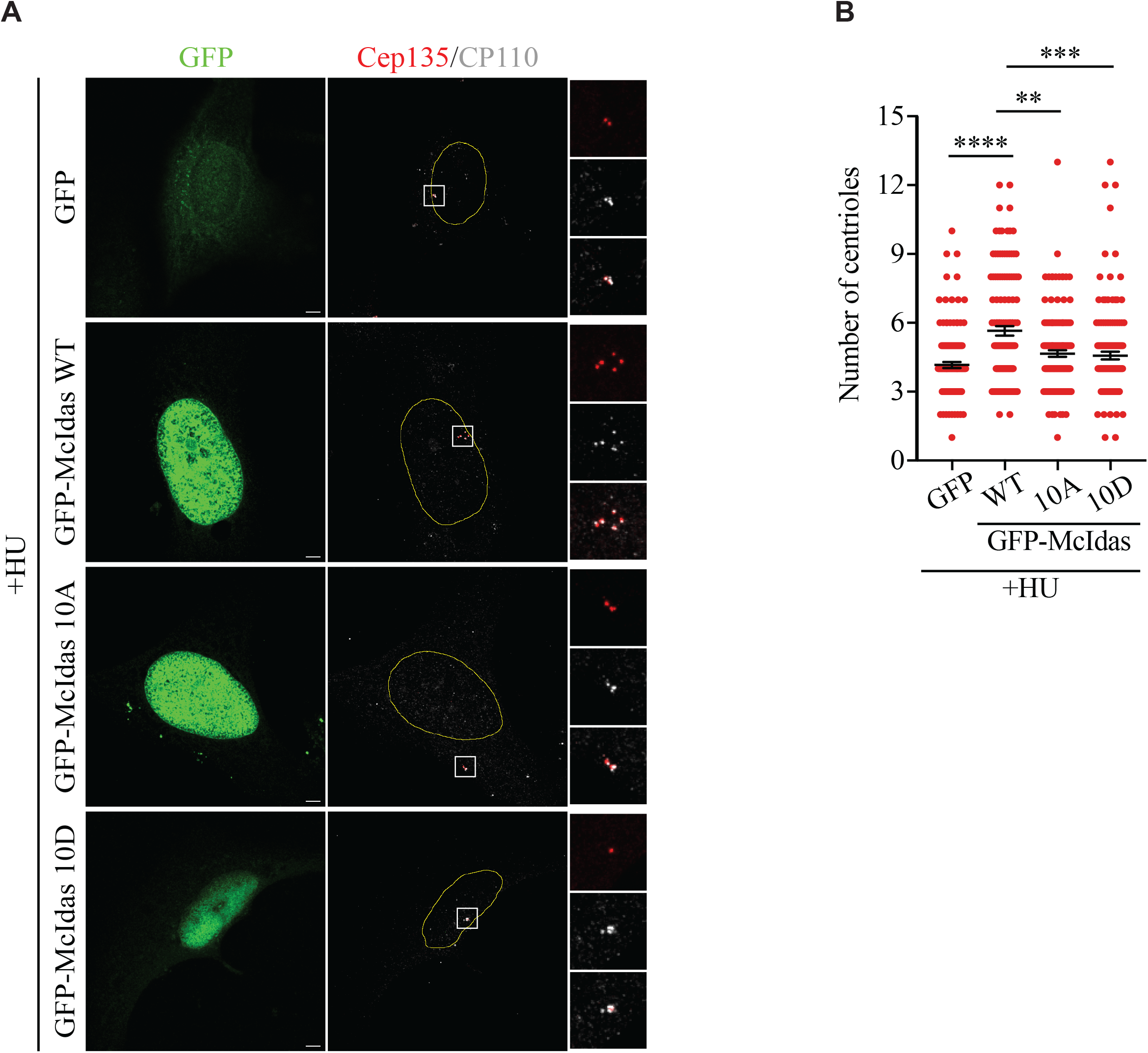
PLK4-specific phosphorylation of McIdas is essential for daughter centriole biogenesis. (A) U2OS cells were treated with 4mM HU and 24 h later were transfected with vectors expressing GFP alone as a control, GFP-McIdas WT, GFP-McIdas 10A (phosphodead mutant) or GFP-McIdas 10D (phospho-mimetic mutant). Cells were fixed 48 h after the transfections and centriole numbers were counted after immunostaining with antibodies against GFP to mark transfected cells, Cep135 and CP110. Inserts corresponds to the higher-magnification images of Cep135 and CP110 stains at centrosomes shown on the right. (B) Quantification of centriole numbers per cell in each condition used at (A). Data are presented as the mean values of two independent experiments and error bars indicate ± SEM. In each experiment at least 100 cells were counted per condition. Nuclei boundaries are marked with lines. Scale bars, 5 μm. *P*-values were calculated using the nonparametric two-tailed Mann-Whitney test. \*\**P* < 0.01, \*\*\**P* < 0.001, \*\*\*\**P* < 0.0001 Abbreviations: HU: hydroxyurea, ns: not significant.

## Discussion

In this study we demonstrate that McIdas has a novel, direct role on centriole duplication. McIdas was originally characterized as a factor implicated in DNA replication initiation (Pefani *et al*., 2011). It was found to be a nuclear protein that interacts with Geminin, affecting Geminin’s affinity for the licensing factor Cdt1. Here, we show that McIdas exhibits an additional centrosomal localization. It is specifically detected at the middle part of centrioles and its centrosomal localization is regulated during the cell cycle. During G1 phase, McIdas is initially present at the mother centriole and then it is recruited at both centrioles. SAS6 is detected at centrioles during S phase, following McIdas recruitment. This is consistent with an early detection of McIdas at centrioles, at the initiation of centriole duplication. Centrosomal McIdas increases as cell cycle progresses and during G2 phase McIdas at mother centrioles is diminished, whereas it is significantly increased at the daughter centrioles (see schematic representations at Figure 2). It would be interesting to investigate whether McIdas differential expression could be linked with processes that take place during G2 phase, including block to reduplication, elongation and maturation of centrioles or separation of centrosomes. McIdas disappears from centrioles at the end of mitosis, during telophase. A previous study showed that McIdas total protein levels display a similar pattern (Pefani *et al*., 2011). Its levels drop during anaphase of mitosis and rise again in early G1 phase. This is reminiscent of the expression pattern followed by other centrosomal regulators, such as STIL and SAS6, which are degraded in late M and early G1 phase by the anaphase-promoting complex/cyclosome associated with Cdh1 (APC/C^Cdh1^). Sequence analysis in McIdas protein revealed a RALL sequence motif in its N-terminal region, which is like the RXXL-type destruction box consensus sequence found in APC/C substrates (Pefani *et al*., 2011). We can speculate that regulation of McIdas expression may also be important to prevent untimely initiation of centriole duplication until the next G1/S phase transition.

Centriole duplication is tightly regulated to avoid reduplication in the same cell cycle. It is known that having excessive centrioles can cause multipolar spindle formation, leading to chromosome mis-segregation during mitosis, and therefore to aneuploidy (Raff and Basto, 2017). Regulation of McIdas expression is important for the maintenance of proper centriole number. McIdas ectopic expression is sufficient to increase centriole numbers in arrested cells, while its loss reduces daughter centriole biogenesis. Our data suggest that McIdas functions in co-operation with PLK4. Immunoprecipitation experiments showed that McIdas is associated with PLK4 and is a novel substrate of this kinase. On the contrary, we were not able to detect binding of McIdas to STIL or SAS6. This could be explained either by low or no affinity of McIdas for these proteins or by the presence of transient protein-protein interactions that we were not able to detect. However, SAS6 recruitment at centrioles is significantly reduced upon McIdas loss. Multiple PLK4-specific phosphorylation sites were identified on McIdas protein at serine residues, through tandem mass spectrometry analysis. The ectopic expression of a McIdas protein that cannot be phosphorylated by PLK4 at the phosphorylation sites mentioned above was not able to increase centriole numbers, as compared to the wild-type protein. This indicates that McIdas phosphorylation by PLK4 is important for its role during centriole duplication. Unexpectedly, the ectopic expression of a phospho-mimetic McIdas mutant also did not increase centriole numbers. This highlights a sophisticated mechanism of McIdas function during centriole duplication cycle, where both its phosphorylation and de-phosphorylation could be important.

Previous work showed that McIdas forms a stable complex with Geminin and prevents Geminin’s binding to Cdt1, suggesting a possible role of McIdas in DNA replication licensing (Pefani *et al*., 2011). Geminin is a negative regulator of DNA replication initiation, inhibiting re-licensing of DNA replication (McGarry and Kirschner, 1998). Moreover, Geminin is localized at centrosome and controls once-per-cell-cycle centrosome duplication (Lu et al., 2009; Tachibana *et al*., 2005; Tachibana and Nigg, 2006). Several other components of the DNA pre-replicative complex, including MCM5, Orc1 and Cdc6, have also been shown to be recruited at centrosomes through interactions with S phase cyclins, controlling centriole duplication (Ferguson and Maller, 2008; Ferguson *et al*., 2010; Hemerly *et al*., 2009; Xu *et al*., 2017). However, the mechanistic understanding of the roles of DNA replication proteins on centriole duplication is still poor. It is intriguing to speculate that McIdas could be a factor that links the centrosome and chromosome cycles, and its function could be mediated through cooperation with other DNA replication licensing proteins.

McIdas was also determined as an important regulator of multiciliogenesis in Xenopus skin and kidney, as well as in mouse airway and brain ependyma (Arbi *et al*., 2016; Boon *et al*., 2014; Kyrousi *et al*., 2015; Stubbs *et al*., 2012). McIdas acts as a transcriptional regulator promoting centriole assembly and ciliogenesis, through the formation of complexes with the transcription factors E2F4/5 and DP1 (Ma *et al*., 2014). A transcriptomic analysis of skin progenitors from Xenopus embryos revealed that McIdas/E2F4/DP1 complex activates not only ciliogenesis-related transcription factors, but also genes essential for centriole assembly, such as PLK4, Cep152, STIL and Deup1 (Ma *et al*., 2014). Recently, Kim *et al*. analyzed McIdas ability to induce multiciliate cell differentiation in primary mouse embryonic fibroblasts (Kim et al., 2018). McIdas overexpression alone caused a significant expansion of centriole number, and this phenotype was significantly enhanced through the co-expression of McIdas with an active form of E2F4. However, a direct role of McIdas on centriole assembly was not examined. Our current findings support an additional centrosomal, direct role of McIdas on centriole biogenesis. Unexpectedly, recent studies revealed an additional role for E2F4 -the binding partner of McIdas-during multicilogenesis, apart from its well-established transcriptional role (Hazan et al., 2021; Mori et al., 2017). E2F4 exhibits a nucleocytoplasmic shift early during multiciliogenesis. This shift was proven to be necessary for organizing centers for nucleation of centrioles in the airway epithelium. Given the centrosomal function of McIdas identified in the present study, it is intriguing to speculate that McIdas could also possess a cytoplasmic role during centriole amplification in multiciliated cells and this could be in co-operation with E2F4.

Multiciliated cells exit the cell cycle and exhibit an S phase-like state in which they massively amplify their centrioles (Lewis and Stracker, 2021; Spassky and Meunier, 2017). But how does the cell choose between duplicating its centrioles once only instead of hundreds of times? The mechanism that controls this balance is still poorly understood. Centriole biogenesis during multiciliate cell differentiation is accomplished through the same regulators that are required for centriole duplication during the cell cycle and Al Jord *et al*. proved that proteins that control cell cycle progression are also essential to drive multiciliated cell differentiation (Al Jord et al., 2017). Very recently, it was suggested that PLK4 protein and its kinase activity are essential for centriole amplification in multiciliated cells (LoMastro et al., 2022). These results corroborate the idea that the initial stages of centriole assembly are conserved between cycling and multiciliated cells. Here we propose that McIdas could be a regulator controlling the decision between two or many centrioles. McIdas expression levels or intricate interactions with different partners could be the responsible mechanisms for McIdas function during a canonical or non-canonical centriole duplication cycle.

In conclusion, our findings identify McIdas as a regulator of centriole duplication cycle in proliferating cells. Given the previously characterized roles of McIdas during the cell cycle and multiciliogenesis, we propose that McIdas has a novel role that links its seemingly unrelated functions.

## Acknowledgements

We thank the Advanced Light Microscopy Facility of the University of Patras for help with microscopy. We are grateful to Dr. Pierre Gönczy for kindly providing SAS6 vectors. We are also thankful to Dr. Constantinos Stathopoulos and Dr. Ilias Skeparnias for their support with the kinase assays and for helpful discussions. We acknowledge the members of Monica Bettencourt-Dias laboratory and especially Catarina Peneda and Dr. Mariana Lince-Faria, as well as Dr. Dafni-Eleftheria Pefani, Dr. Ioannis Loukas, and the members of Zoi Lygerou & Stavros Taraviras laboratories for insightful discussions regarding this work.

This work was supported by: the Hellenic Foundation of Research and Innovation (HFRI) and General Secretariat for Research and Innovation (GSRT) under grant agreement No 1302 to M.A., the project “BioimagingGR” (MIS 5002755) to Z.L. and the project “INSPIRED – The National Research Infrastructures on Integrated Structural Biology, Drug Screening Efforts and Drug target functional characterization” (MIS 5002550) to G.A.S., which are implemented under the Action “Reinforcement of the Research and Innovation Infrastructure”, funded by the Operational Programme “Competitiveness, Enterpreneurship and Innovation” (NSRF 2014-2020) and co-financed by Greece and the European Union (European Regional Development Fund).

## Author contributions

Conceptualization, M.A., Z.L.; Methodology, M.A., S.Z., O.K., C.G.V., A.C.T., N.N.G.; Investigation (microscopy, molecular biology and biochemical analysis), M.A., M.S., S.B., S.Z., S.T., A.C.T., N.N.G.; Investigation (LC-MS/MS analysis, data processing), O.K., C.G.V; Resources, G.A.S., M.M., M.B.D., S.T., Z.L.; Writing – Original Draft, M.A.; Writing – Review & Editing, Z.L., M.B.D.; Visualization, M.A., O.K.; Supervision, M.A., S.T., Z.L.; Project administration, M.A., Z.L.; Funding acquisition, M.A., Z.L.

## Materials and Methods

### Plasmids

The control GFP plasmid (pLVDest-GFP) used in this study was previously described (Arbi *et al*., 2016). For the construction of a GFP-tagged mouse McIdas plasmid, the full-length mouse McIdas cDNA (Pefani *et al*., 2011) with an N-terminal GFP tag was cloned into the KpnI/XhoI restriction sites of the pENTR1AminusCmR vector. For the construction of a human GFP-tagged McIdas vector, the full-length cDNA of human McIdas was obtained by GenScript (OHu00715) and cloned with an N-terminal GFP tag into the EcoRI/XhoI restriction sites of the pENTR1AminusCmR vector. The phospho-dead and phospho-mimetic mutant forms of McIdas were obtained by GenScript. In short, 10 serine residues on McIdas protein (S55, S58, S86, S87, S167, S246, S271, S333, S338 and S340) were mutated either to alanine (phospho-dead mutant, McIdas 10A) or to aspartic acid (phospho-mimetic mutant, McIdas 10D) residues. Each McIdas mutant was cloned with an N-terminal GFP tag into the EcoRI/XhoI restriction sites of the pENTR1AminusCmR vector to generate the GFP-McIdas 10A and GFP-McIdas 10D constructs. The GFP-McIdas 10A construct was used to generate the partial phospho-dead McIdas mutants used here (4A_N, 3A_cc & 3A_C). Briefly, to generate the GFP-McIdas 4A_N, GFP-McIdas 3A_cc and GFP-McIdas 3A_C mutants, subcloning from the GFP-McIdas 10A plasmid to the GFP-McIdas WT plasmid was performed by using the SacII/MscI, MscI/KasI and KasI/XhoI restriction sites, respectively. Using the Gateway LR clonase II enzyme mix (Invitrogen, 11791), LR recombination reactions were performed between the attL-containing entry clones and attR-containing destination pLVDest-CAG vectors to produce the following final expression vectors: pLVDest-GFP-McIdas (mouse), pLVDest-GFP-McIdas (human), pLVDest-GFP-McIdas 10A, pLVDest-GFP-McIdas 10D, pLVDest-GFP-McIdas 4A_N, pLVDest-GFP-McIdas 3A_cc and pLVDest-GFP-McIdas 3A_C.

Plasmid vectors used in the co-immunoprecipitation experiments are listed below: pcDNA3.1-McIdas-HA and pcDNA3.1-HA control vectors (Pefani *et al*., 2011), pCMV-HA-STIL (Ohta *et al*., 2014), pEBTet-HsSAS6-GFP and pEBTet-HsSAS6_SNAP-FLAG (1 μg/ml tetracycline was used for efficient induction, provided by Dr. Pierre Gonczy, Keller et al 2014).

### Cell culture and treatments

U2OS (ATCC, HTB-96), HeLa (ATCC, CCL-2) and 293T (ATCC, CRL-3216) cells were cultured in DMEM (Gibco) supplemented with 10% fetal bovine serum in 5% CO_2_ at 37 °C. hTERT-RPE1 cells (ATCC, CRL-4000) were cultured in DMEM/F12 (Sigma-Aldrich) with 10% fetal bovine serum in 5% CO_2_ at 37 °C. U2OS cells expressing myc-PLK4 under a tetracycline-dependent promoter (Kleylein-Sohn et al., 2007) were grown in DMEM supplemented with 10% tetracycline-free fetal bovine serum (PAN-Biotech) in 5% CO_2_ at 37 °C. For an efficient myc-PLK4 overexpression, cells were grown in medium containing 100 ng/ml tetracycline (Sigma-Aldrich).

For the construction of the stable cell lines, HeLa cells were infected with lentiviral particles expressing GFP or GFP-McIdas. A second-generation packaging system was used to produce lentiviral particles, as described previously (Arbi *et al*., 2016). Briefly, 293T cells were transiently co-transfected with the expression vector carrying either the GFP (pLVDest-GFP) or the GFP-MCIDAS gene (pLVDest-GFP-McIdas) and the two helper plasmids psPAX2 (packaging vector, Addgene 12260) and pMD2.G (envelope vector, Addgene 12259), using Turbofect (Fermentas) as transfection reagent. The supernatant was harvested 48 h after the transfection and filtered. HeLa cells were infected with the lentiviral preparations containing 5 μg/ml hexadimethrine bromide (polybrene, Sigma-Aldrich). Limiting dilution and single colony picking were used to generate the HeLa stable cell lines. To inhibit proteasome activity, Hela cells stably expressing GFP or GFP-McIdas were treated with MG132 (100 μM, Sigma-Aldrich) for 4 h.

### HU-induced centriole amplification assay and transfections

U2OS cells were cultured in a 6-well plate at 80% confluency and treated with 4 mM HU. 24 h later cells were transfected, using polyethylenimine (PEI, 3 μg/ml, Polysciences, 23966), with 1 μg of the indicated plasmids. Cells were fixed and stained 48 h post transfection.

### RNA interference

For McIdas RNAi experiments the following STEALTH siRNA oligos (McIdas siRNA_1; Invitrogen), 5’-CCACCAAACGGAAGCAGACTTCAAT-3’ and 5’-GAGACGCGCTTGTTGAGAATAATCA-3’, were used as described previously (Pefani *et al*., 2011). A luciferase siRNA was used as a control: 5’-CGTACGCGGAATACTTCGA-3’. A second siRNA oligo (McIdas siRNA_2) was also used for McIdas depletion: 5’-CAACTGCACGTGACATTGATT-3’ (s51151, Silencer® Select siRNA, Thermo Fisher Scientific). The Silencer® Select No. 1 siRNA (4390843) was used as a negative control (Thermo Fisher Scientific). U2OS or U2OS-mycPLK4 cells were seeded onto coverslips at 80% confluency, in a 6-well plate and transfected with the indicated siRNA oligos, using RNAiMAX (Life Technologies) according to the manufacturer’s instructions. A second treatment with the siRNA oligos followed 24 h later. McIdas siRNA_1 or luciferase siRNA oligos were used at a final concentration of 80 nM, whereas McIdas siRNA_2 or control siRNA oligos were used at a final concentration of 40 nM. For HU-induced centriole amplification, 5 h after the second transfection, cells were treated with 4 mM HU. For PLK4-induced centriole amplification, U2OS-mycPLK4 cells, 5 h after the second transfection, were cultured in medium containing 100 ng/ml tetracycline. For the detection of S phase cells, EdU (10 μM, Invitrogen) was added in the culture medium for 1 h before fixation. Cells were analyzed either with qPCR or western blotting for the quantification of McIdas mRNA and protein levels or fixed and stained, 48 h after the second transfection.

### Immunofluorescence and expansion microscopy (ExM)

Cells were grown on coverslips, washed twice with PBS 1x, and fixed with precooled (-20 °C) methanol for 10 min at -20 °C. For the detection of GFP-tagged McIdas at centrosomes pre-extraction was performed before methanol fixation. Briefly, cells were treated with CSK buffer (10 mM PIPES pH 7.9, 300 mM sucrose, 100 mM NaCl and 3 mM MgCl_2_) for 1 min at room temperature, followed by incubation with CSK + Triton X-100 buffer for 1 min at room temperature (0.5 % Triton X-100, 1 mM PMSF, 1x protease inhibitor cocktail in CSK buffer). To visualize centrioles with acetylated α-tubulin for ExM, cells were incubated on ice for 30 min to depolymerize cytoplasmic microtubules before fixation. Staining of S phase cells was performed with the Click-iT Alexa Fluor 647 Imaging Kit (Invitrogen) after fixation, according to the manufacturer’s instructions. Cells were subsequently blocked in solution containing 10% FBS and 3% BSA, in 1x PBS, for 1 h, followed by incubation with primary antibodies in blocking solution at 4 °C, overnight. The primary antibodies used were as follows: rabbit anti-hMcIdas (1:250) (Pefani *et al*., 2011), mouse anti-Centrin (1:500-1:1000, Merck Millipore), chicken anti-GFP (1:1000, Aves Labs), rat anti-Cep135 (1:500, Monica Bettencourt-Dias lab, rat Cep135 antibody produced against human CEP135 by Metabion. For immunization, a region of CEP135, aa 270-371, was expressed in *E. coli* as a fusion protein with a N-terminal 10x His tag), goat anti-Cep164 (1:500, Santa Cruz Biotechnology), mouse anti-SAS6 (1:500, Santa Cruz Biotechnology), rabbit anti-CP110 ((1:250, Jiang et al., 2012), rabbit anti-PLK4 (1:500, Monica Bettencourt-Dias lab, rabbit PLK4 antibody was produced against the c-terminal region, aa 510-970, of human PLK4 and purified using the membrane-bound c-terminal region of PLK4). After three washes with PBS 1x-Tween 0.1 %, Alexa Fluor-labeled secondary antibodies (Invitrogen) were used at 1:500 in blocking solution for 1h at room temperature: Alexa Fluor 488 goat anti-Rabbit, Alexa Fluor 568 goat anti-Mouse, Alexa Fluor 488 goat anti-Chicken, Alexa Fluor 488 goat anti-Rat, Alexa Fluor 568 donkey anti-Goat and Alexa Fluor 647 donkey anti-Rabbit. DNA was stained with Hoechst 33258 (1:1000, Sigma). Coverslips were mounted in Mowiol 4-88 (Calbiochem) or further processed for ExM (Asano et al., 2018).

For ExM, cells were incubated with 0.1 mg/ml Acryloyl-X (Thermo Fisher Scientific) in PBS 1x at room temperature, overnight. Cells were washed twice with PBS 1x for 15 min and incubated in monomer solution (900 mM Sodium Acrylate, 2.6% Acrylamide 37.5:1, 0.15% N, N’-Methylenebisacrylamide, 2M NaCl, PBS 1x) containing 4-HT (0.01 %), APS (0.2 %) and TEMED (0.2 %) in a 47:1:1:1 ratio, for 2 h at 37 °C. After complete polymerization, gels were digested with 8 U/ml Proteinase K (Thermo Fisher Scientific) in a solution containing 50 mM Tris-HCl pH 8, 1 mM EDTA, 1 M NaCl and 0.5% Triton X-100, for 1 h at 37 °C. Gels were expanded in milli-Q water (3 × 20 min) and imaged on poly-L-lysine-coated μ-slide 2 well glass bottom (Ibidi). Expansion factor was approximately 4x. The primary and secondary antibodies used for the expanded samples were as follows: rabbit anti-hMcIdas (1:100) (Pefani *et al*., 2011), mouse anti-acetylated α-tubulin (1:250, Sigma), Alexa Fluor 488 goat anti-Rabbit (1:250, Invitrogen) and Alexa Fluor 568 goat anti-Mouse (1:250, Invitrogen).

Images were taken using a Leica TCS SP5 confocal system. Image acquisition was done with a 63x NA 1.4 oil immersion lens, XY kept at 512 × 512 or 1024 × 1024 pixels with a bidirectional scanning speed of 400 Hz. Z-stacks were acquired with a 0.2 μm step. For the expanded samples we acquired the images in a frame-by-frame manner to minimize gel drift. Digital images were processed with ImageJ software (Rueden et al., 2017). Image intensity quantifications were performed with the use of a custom ImageJ macro on the SUM projections of Z-Stack. Multichannel intensity profiles, as a function of distance on a line connecting the local maximal intensity points marking the centrioles, were calculated with the use of a custom ImageJ macro.

**Co-immunoprecipitation and western blot analysis**

Immunoprecipitation was performed as described previously (Lalioti et al., 2019). U2OS cells expressing myc-PLK4 under a tetracycline-dependent promoter or 293T cells were transiently transfected in 10 cm plates, using polyethylenimine (PEI), with a total of 6 μg of the indicated plasmids. For myc-PLK4 overexpression, 100 ng/ml tetracycline (Sigma-Aldrich) was added to the growth medium, 5 h post-transfection. Cells were collected 48 h after transfection, washed twice with ice cold PBS 1x and incubated for 10 min in lysis buffer (50 mM Tris-HCl pH 8.2, 150 mM NaCl, 5 mM EDTA, 5 mM MgCl_2_, 0.5% Triton X-100) supplemented with fresh protease and phosphatase inhibitors (1 mM PMSF, 0.1 mM Na_3_VO_4_ and 1x PIC - Roche). Cell lysates were passed through a 1 ml syringe for mechanical disruption and centrifuged for 10 min at 13.000 rpm. In parallel, for each IP reaction, 50 μl of Protein A agarose suspension (Merck Millipore, IP02) was incubated with 1.5 μg of the indicated antibodies, for 2 h on a rotating wheel at 4 °C. Beads were washed three times with lysis buffer and incubated with 1-2 mg of total protein from whole-cell lysates, for 3 h on a rotating wheel at 4 °C. Beads were washed again with lysis buffer and proteins were eluted in Laemmli buffer by boiling for 5 min. Proteins were analyzed by SDS-PAGE and transferred onto PVDF membrane (Merck Millipore). The membranes were washed with PBS 1x containing 0.1% Tween-20 (PBS-T) and blocked in 5% milk (in PBS-T) for 1 h, at room temperature, followed by incubation with primary antibodies at room temperature, overnight. The membranes were incubated with HRP-conjugated secondary antibodies for 1 h, at room temperature and exposed either to X-ray film (Santa Cruz Biotechnology) or to the ChemiDoc Imaging System (Biorad) after incubation with the Clarity ECL (Biorad). The antibodies used in immunoprecipitation were as follows: mouse anti-GFP (clones 7.1 & 13.1; 11814460001, Roche) and mouse anti-HA (F-7 clone; sc-7392, Santa Cruz Biotechnology). The primary and secondary antibodies used in western blot analysis were: mouse anti-GFP (1:1000, Roche), mouse anti-PLK4 (1:500, Millipore), mouse anti-HA (1:1000, Santa Cruz Biotechnology), mouse anti-Flag (1:1500, Sigma-Aldrich), goat anti-Rabbit IgG-HRP (1:1000-1:3000, Merck Millipore) and goat anti-Mouse IgG-HRP (1:1000-1:3000, Merck Millipore).

### Protein expression and purification

For *in vitro* kinase assays three truncated McIdas recombinant proteins were used: an N-terminal fragment (1-179 aa), a fragment containing the central coiled-coil domain (101-284 aa) and a fragment containing the conserved C-terminal region of McIdas (314-385 aa). McIdas recombinant proteins containing the N-terminal region (similar to McIdas 1-127 aa) and the central coiled-coil domain (101-284 aa) have been described previously (Pefani *et al*., 2011). The C-terminal region of McIdas (314-385 aa) was amplified from a synthetic and codon optimized gene purchased from GenScript, and cloned into pET20b(+) vector. The used primer sequences were: FW 5’-GGAATTCCATATGACCCGTCCGGGTAACC-3’, RV 5’-CCGGCTCGAGGCTCGGCACCCAACGGAACTT-3’. The obtained construct was verified by DNA sequencing. The final derived polypeptide is expressed fused with a C-terminal His_6_-tag.

For the expression of the C-terminal region of McIdas (314-385 aa), *Escherichia coli* cells (BL21 (DE3) Singles™ Competent Cells – Novagen) were transformed with the vector. Cells were grown in LB medium containing ampicillin till an O.D. value 0.6-0.8. Expression was induced with 1 mM isopropyl β-D-1-thiogalactopyranoside (IPTG) for 4 h at 37 °C. Cells were harvested and resuspended in buffer containing 50 mM Tris-HCl pH 8, 300 mM NaCl, 0.1% Triton X-100. After two repeated cycles of sonication and centrifugation the derived pellet was resuspended in 50 mM Tris-HCl pH 8, 500 mM NaCl, 10 mM Imidazole, 2 mM β-mercaptoethanol and 6 M urea followed by 2 h agitation and centrifugation at 14.000 rpm. The supernatant was loaded onto a HisTrap column (GE Healthcare) and the C-terminal region of McIdas was purified in presence of urea followed by buffer exchange in 50 mM Tris-HCl pH 8, 300 mM NaCl and 2 mM β-mercaptoethanol. The desired concentration was achieved using Amicon® Ultra centrifugal filter units (3 kDa cutoff – Merck Millipore).

### Kinase assays

293T cells were transiently transfected in 10 cm plates, using polyethylenimine (PEI), with 6 μg of the indicated plasmids. Cells were collected 48 h after transfection, washed twice with ice cold PBS 1x and incubated for 10 min in lysis buffer without phosphatase inhibitors (50 mM Tris-HCl pH 8.2, 150 mM NaCl, 0.5% Triton X-100) supplemented with fresh protease inhibitors (1 mM PMSF and 1x PIC - Roche). Cell lysates were passed through a 1 ml syringe for mechanical disruption and centrifuged for 10 min at 13.000 rpm. To release phosphate groups from phosphorylated residues in proteins, cell lysates were subjected to phosphatase treatment for 30 min at 30 °C, according to the manufacturer’s instructions (Lambda protein phosphatase, Biolabs). In parallel, for each IP reaction 50 μl of Protein A agarose suspension (Merck Millipore, IP02) was incubated with 1.5 μg of GFP antibody (mouse anti-GFP, clones 7.1 & 13.1; 11814460001, Roche), for 2 h on a rotating wheel at 4 °C. Beads were washed three times with lysis buffer without phosphatase inhibitors and incubated with 1-2 mg of total protein from whole-cell lysates, for 3 h on a rotating wheel at 4 °C. Beads were washed three times with lysis buffer (50 mM Tris-HCl pH 8.2, 150 mM NaCl, 5 mM EDTA, 5 mM MgCl_2_, 0.5% Triton X-100) supplemented with fresh protease and phosphatase inhibitors (1 mM PMSF, 0.1 mM Na_3_VO_4_ and 1x PIC - Roche), followed by three washes with kinase buffer (25 mM Tris-HCl pH 7.5, 25 mM NaCl, 10 mM MgCl_2_ and 1 mM DTT). Each sample was incubated with 500 ng recombinant Xenopus PLK4 (PLX4, provided by Dr. Monica Bettencourt-Dias) and 100 μM [γ-^32^P] ATP for 30 min at 30 °C. Samples were analyzed by SDS-PAGE followed by autoradiography. Western blot analysis was performed in the 10-20% of each sample, as well as in the total cell extracts used for the kinase assays. The primary and secondary antibodies used in this analysis were: mouse anti-GFP (1:1000, Roche), rabbit anti-PLK4 (1:500), mouse anti-α-tubulin (1:6000, Sigma-Aldrich), goat anti-Rabbit IgG-HRP (1:1000-1:3000, Merck Millipore) and goat anti-Mouse IgG-HRP (1:1000-1:3000, Merck Millipore). The kinase reactions using recombinant McIdas were performed as follows: 1 μg of each McIdas recombinant protein was incubated with 500 ng recombinant Xenopus PLK4 (PLX4) in the presence of 100 μM ^32^P-radiolabeled ATP for 30 min at 30 °C. An irrelevant recombinant protein was used as a negative control. Samples were analyzed by SDS-PAGE followed by autoradiography and Coomassie staining or subjected to tandem mass spectrometry analysis.

### LC-MS/MS Analysis and Data Processing

Recombinant proteins were diluted 1:1 in 100 mM Tris-HCl (pH 8), containing 10 mM TCEP and 40 mM 2-Chloroacetamide to the final concentrations of 10 mM and 40 mM, respectively. Trypsin alone or in combination with AspN was then added to samples which were subsequently incubated for 4 h at 37 °C with agitation (1,500 rpm). Next day, peptides were loaded onto SDB-RPS StageTips after acidifying with 10% TFA to ∼1% and desalted as described previously (Kulak et al., 2014). Briefly, the StageTips were centrifuged at 1,000 g for washing with 2% ACN/0.2% TFA twice and at 500 g for elution with 80% ACN/0.1% TFA. The eluate was evaporated to dryness using a vacuum centrifuge and peptides were resuspended in MS loading buffer (2% ACN/0.2% TFA). Equal amount of peptides was subjected to LC-MS/MS analysis.

Peptides were separated on a 50 cm reversed-phase column (75 μm inner diameter, packed in-house with ReproSil-Pur C18-AQ 1.9 μm resin [Dr. Maisch GmbH]) with a binary buffer system of buffer A (0.1% formic acid (FA)) and buffer B (80% acetonitrile plus 0.1% FA) over 60 min gradient (steps: (1) 5%–30% of buffer B for 35 min, (2) 30%–65% for 5 min, (3) 65%-95% for 5 min and (4) washout for 15 min) using the EASY-nano LC 1200 system (Thermo Fisher Scientific) with a flow rate of 300 nL/min. Column temperature was maintained at 60°C. The nano LC system was coupled to Orbitrap Exploris 480 mass spectrometer (Thermo Fisher Scientific). The instrument is operated in Top12 DDA mode. We acquired full scans (300–1,650 m/z, maximum injection time 25 ms, resolution 60,000 at 200 m/z, charges included 2-5 and dynamic exclusion of 30 ms) at a target of 3e6 ions. The 12 most intense ions were isolated and fragmented with higher-energy collisional dissociation (HCD) (target 1e5 ions, maximum injection time 28 ms, isolation window 1.4 m/z, NCE 28%) and detected in the Orbitrap (resolution 15,000 at 200 m/z).

Raw MS files were processed within the MaxQuant environment (version 1.6.7.0) with the integrated Andromeda search engine with FDR < 0.01 at the protein and peptide levels (Cox et al., 2014; Cox and Mann, 2008; Cox et al., 2011). We included methionine (M) oxidation and acetylation (protein N-term) as variable and carbamidomethyl (C) as fixed modifications in the search. We allowed up to 2 missed cleavages for tryptic and AspN digestion and considered peptides with at least six amino acids for identification. ‘‘Match between runs’’ was enabled with a matching time window of 0.7 min to allow the quantification of MS1 features which were not identified in each single measurement. Peptides and proteins were identified using a UniProt FASTA database from *Homo sapiens* (2015) containing 21,051 entries.

The mass spectrometry proteomics data have been deposited to the ProteomeXchange Consortium via the PRIDE (Perez-Riverol et al., 2022) partner repository with the dataset identifier PXD037043.

### Phosphatase treatment

293T cells were transfected with vectors expressing GFP-McIdas WT or the mutant forms, GFP-McIdas 10A and GFP-McIdas 10D. Cell lysates were collected and incubated with lambda protein phosphatase for 30 min at 30 °C, according to the manufacturer’s instructions (Lambda protein phosphatase, Biolabs). Western blot analysis was followed with an antibody against GFP (1:1000, mouse anti-GFP, Roche) and goat anti-Mouse IgG-HRP (1:3000, Merck Millipore).

### RNA purification and real-time PCR

To assess McIdas mRNA levels, total RNA after McIdas RNAi was isolated from U2OS cells using the Nucleospin RNA II kit (Macherey-Nagel). 1 μg RNA was converted to cDNA using M-MLV transcriptase (Invitrogen). McIdas mRNA levels were assessed by quantitative real-time PCR (StepOne, Applied Biosystems), using the Kapa SYBR Fast qPCR kit (Applied Biosystems), according to the manufacturer’s instructions. YWHAZ mRNA expression levels were used for normalization. qPCR data analysis was performed with the REST-MCS beta software. Primer sequences used were as follows: hMcIdas FW 5’-GACGCGCTTGTTGAGAATAA-3’, hMcIdas RV 5’-CACGTTCCGCTCCTTGAG-3’, YWHAZ FW 5’-GATCCCCAATGCTTCACAAG-3’ and YWHAZ RV 5’-TGCTTGTTGTGACTGATCGAC-3’.

## Supplementary Figures

**Supplementary Figure 1. McIdas localizes at centrosomes**

(A, B) U2OS (A) and hTERT-RPE1 (B) cells were transfected with vectors expressing either GFP-tagged McIdas or GFP alone as a control. Cells were pre-extracted and immunostained with antibodies against GFP to mark the transfected cells, McIdas and Centrin (distal lumen centriole marker). Inserts show higher magnification images of the indicated stains at centrosomes. Arrows correspond to the position of centrosome and asterisks show increased accumulation of Centrin upon GFP-McIdas overexpression. (C) HeLa cells were generated to stably express a GFP-tagged McIdas protein or GFP alone as a control and treated with the proteasome inhibitor MG132 for 4 h. Cells were fixed and immunostained with antibodies against GFP, endogenously expressed McIdas and Cep135 (proximal centriole marker). Inserts show higher magnification images of McIdas stain at centrosomes. (D-G) Two different McIdas siRNA oligos (1 and 2) were used to verify the specificity of McIdas localization at centrosomes. U2OS cells were transfected with luciferase or McIdas siRNA_1 (D, E) and control or McIdas siRNA_2 (F, G). Cells were collected and McIdas mRNA levels were assessed by qPCR (D, F) or were immunostained with antibodies against McIdas to count McIdas fluorescence intensity at centrosomes (E, G). Data from a representative experiment were shown. At (D) and (F) samples were analyzed in duplicates. For the quantifications of McIdas centrosomal levels (E, G) at least 40 centrosomes were counted per condition. Two independent experiments were conducted for all data shown. Error bars indicate ± SEM. *P*-values in E and G were calculated by the nonparametric two-tailed Mann-Whitney test. \**P* < 0.1, \*\**P* < 0.01.

DNA was stained with Hoechst. Scale bars, 5 μm.

Abbreviations: ns: not significant.

**Supplementary Figure 2. McIdas contributes to daughter centriole biogenesis in cells**

(A) McIdas siRNA_1 or control siRNA oligos were transfected in U2OS cells. 24 h later, after a second transfection with the indicated siRNAs, cells were treated with 4mM HU for another 48 h. McIdas mRNA levels were assessed by qPCR. (B, C) U2OS cells expressing myc-PLK4 under a tetracycline-dependent promoter were transfected with McIdas_2 or control siRNA oligos. A second transfection with the indicated siRNAs followed 24 h later and then cells treated with tetracycline for an efficient myc-PLK4 overexpression. McIdas protein (B) and mRNA (C) levels were assessed by western blot and qPCR, respectively. (D) U2OS cells expressing myc-PLK4 under a tetracycline-dependent promoter were transfected with McIdas_2 or control siRNA oligos. A second transfection with the indicated siRNAs followed 24 h later and then cells treated with tetracycline for an efficient myc-PLK4 overexpression. Cells were fixed and immunostained for Centrin to count centriole numbers. (E) Representative quantification of centriole numbers in control and McIdas-depleted cells, overexpressing PLK4. At least two independent experiments have been performed. In each experiment more than 100 cells were counted per condition. (F, G) U2OS cells expressing myc-PLK4 under a tetracycline-dependent promoter were treated with tetracycline or DMSO as a control and immunostained with antibodies against McIdas and PLK4 (F) or SAS6 (G). (H) U2OS cells were transfected with McIdas_2 or control siRNA oligos and McIdas mRNA levels were assessed by qPCR.

Nuclei boundaries are marked with lines. DNA was stained with Hoechst. Scale bars, 5 μm. Inserts corresponds to higher-magnification images of the indicated markers at centrosomes, shown on the right.

*P*-values in A, C and H were calculated using two-tailed Student’s t-test. *P*-value in E was calculated by the nonparametric two-tailed Mann-Whitney test. \*\**P* < 0.01, \*\*\**P* < 0.001, \*\*\*\**P* < 0.0001

Abbreviations: HU: hydrxyurea, Tet: tetracycline, ns: not significant.

**Supplementary Figure 3. McIdas is not able to interact with STIL and SAS6**

(A, B) 293T cells were co-transfected with vectors expressing HA-STIL and GFP-McIdas or GFP alone as a control. Cell extracts were collected and immunoprecipitation was performed either against GFP (A) or against HA (B). STIL was not detected in McIdas immunoprecipitates or *vice versa*. (C) 293T cells were co-transfected with vectors expressing SAS6-GFP under a tetracycline-dependent promoter and McIdas-HA or HA alone as a control. Cell extracts were collected and immunoprecipitation was performed against HA. (D) 293T cells were co-transfected with vectors expressing SAS6-SNAP-Flag under a tetracycline-dependent promoter and GFP-McIdas or GFP alone as a control. Cell extracts were collected and immunoprecipitation was performed against GFP.

Asterisks indicate the specific bands in each experiment.

Abbreviations: TCL: total cell extract, IP: immunoprecipitation.

**Supplementary Figure 4. PLK4-dependent phosphorylation of McIdas is enriched on its coiled coil-containing region**

(A) 293T cells were transfected with vectors expressing GFP-McIdas WT protein or the mutant forms 10A and 10D. Cell lysates were collected and incubated with lambda protein phosphatase. Western blot analysis was followed with an antibody against GFP. (B) Schematic representation of the PLK4-phosphorylated serine residues identified on McIdas protein and the partial McIdas phospho-dead mutant forms that have been generated. (C) *In vitro* kinase assays were performed using IP control or GFP-McIdas WT or the partial phospho-dead GFP-McIdas mutants, as indicated. Each IP was incubated with recombinant Xenopus PLK4 (PLX4) and [γ-^32^P] ATP. Samples were analyzed by autoradiography. Western blot analysis was performed in the 20% of each sample by using antibodies against PLK4 and GFP to monitor loaded proteins. (D) Western blot analysis of total cell extracts used at (C) for the *in vitro* kinase assays was performed by using antibodies against GFP and tubulin. (E) Representative quantification of phosphorylation of McIdas WT or partial mutant forms used at (C), calculated by Image Lab software. Values indicate phosphorylation signal intensity relative to McIdas WT (set as 1). At least 2 independent experiments have been performed.

