## Supplementary figures and images for "McIdas localizes at centrioles and controls centriole numbers through PLK4-dependent phosphorylation"

### Supplementary Figure 1

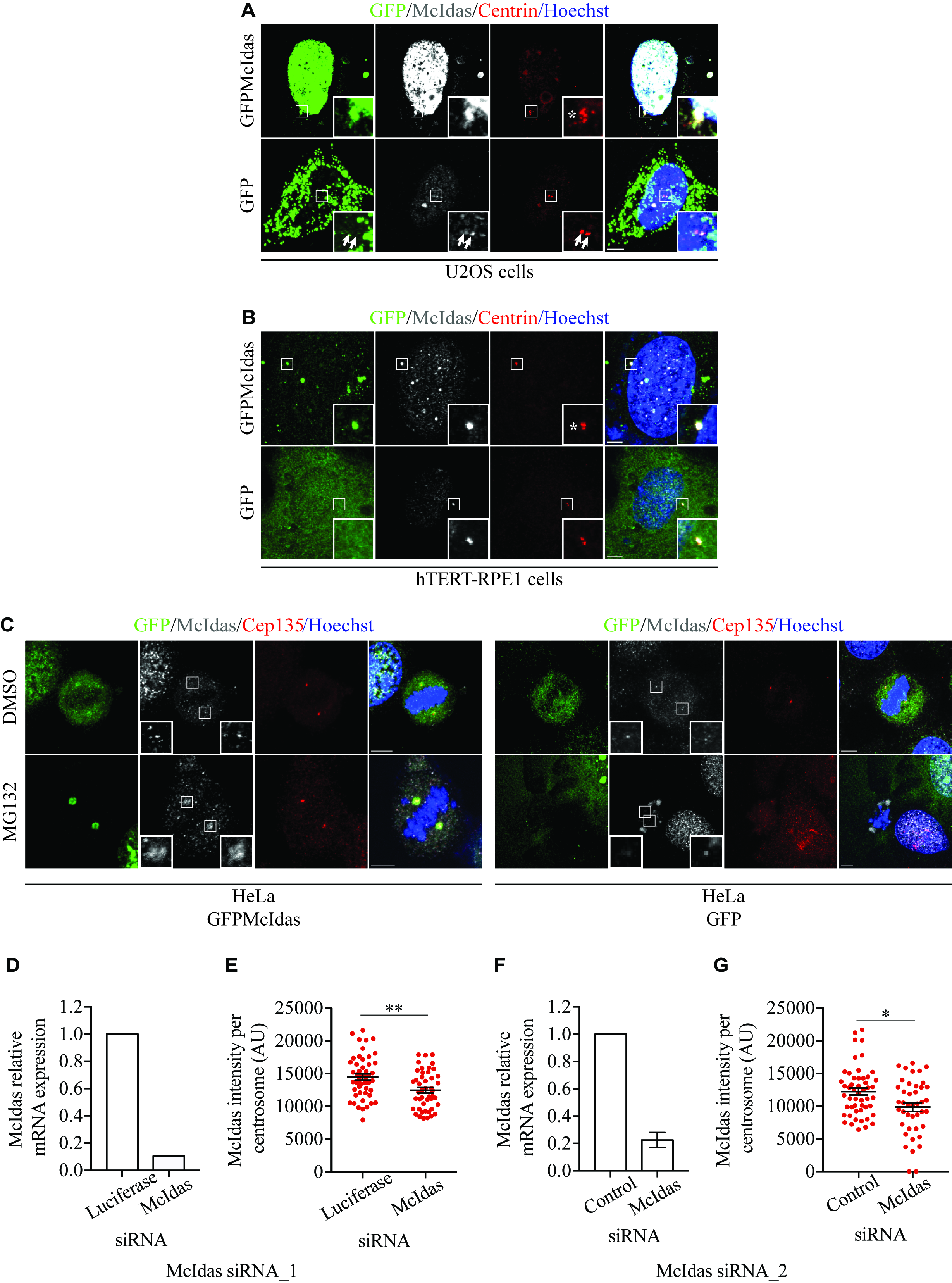

### Supplementary Figure 2

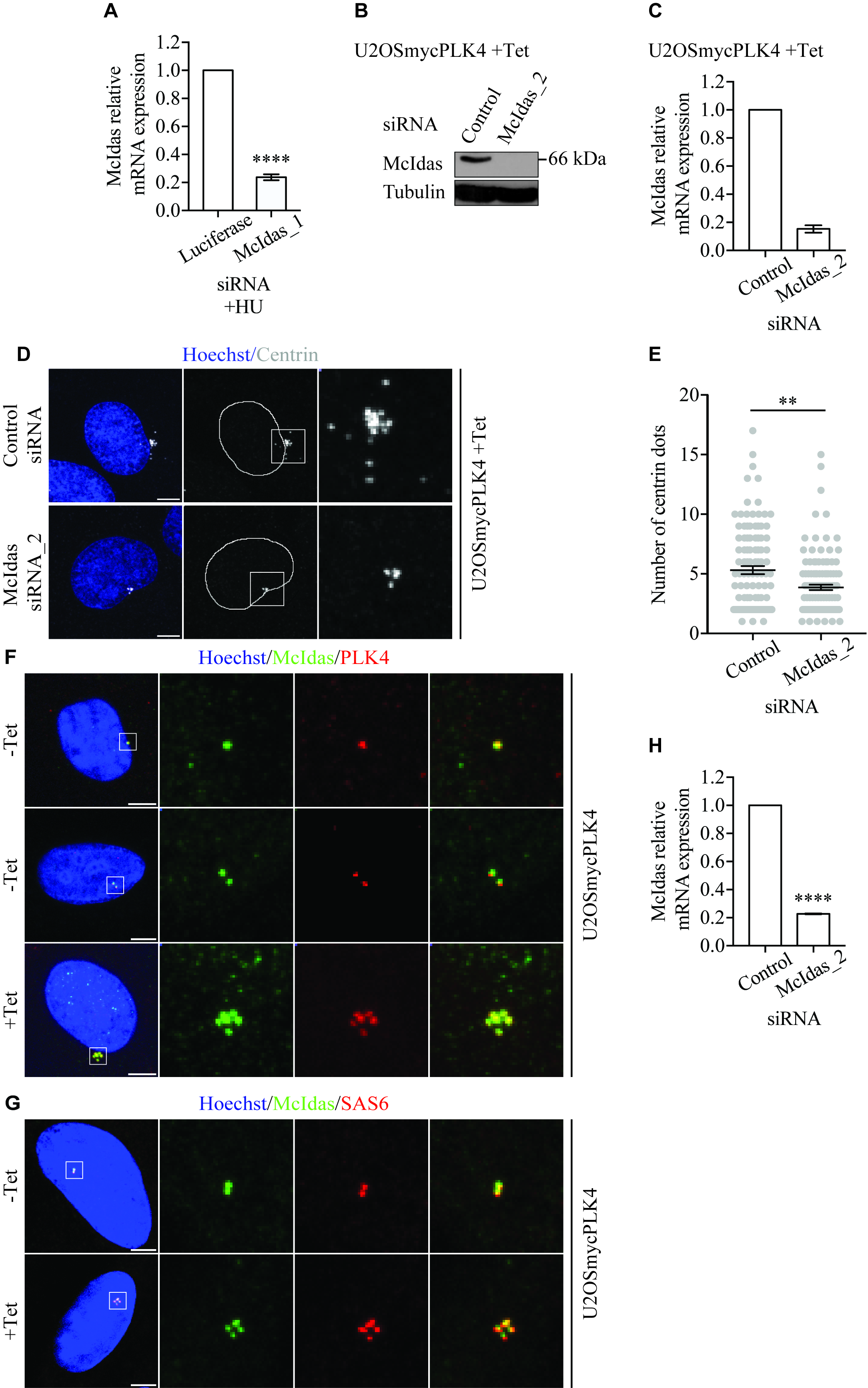

### Supplementary Figure 3

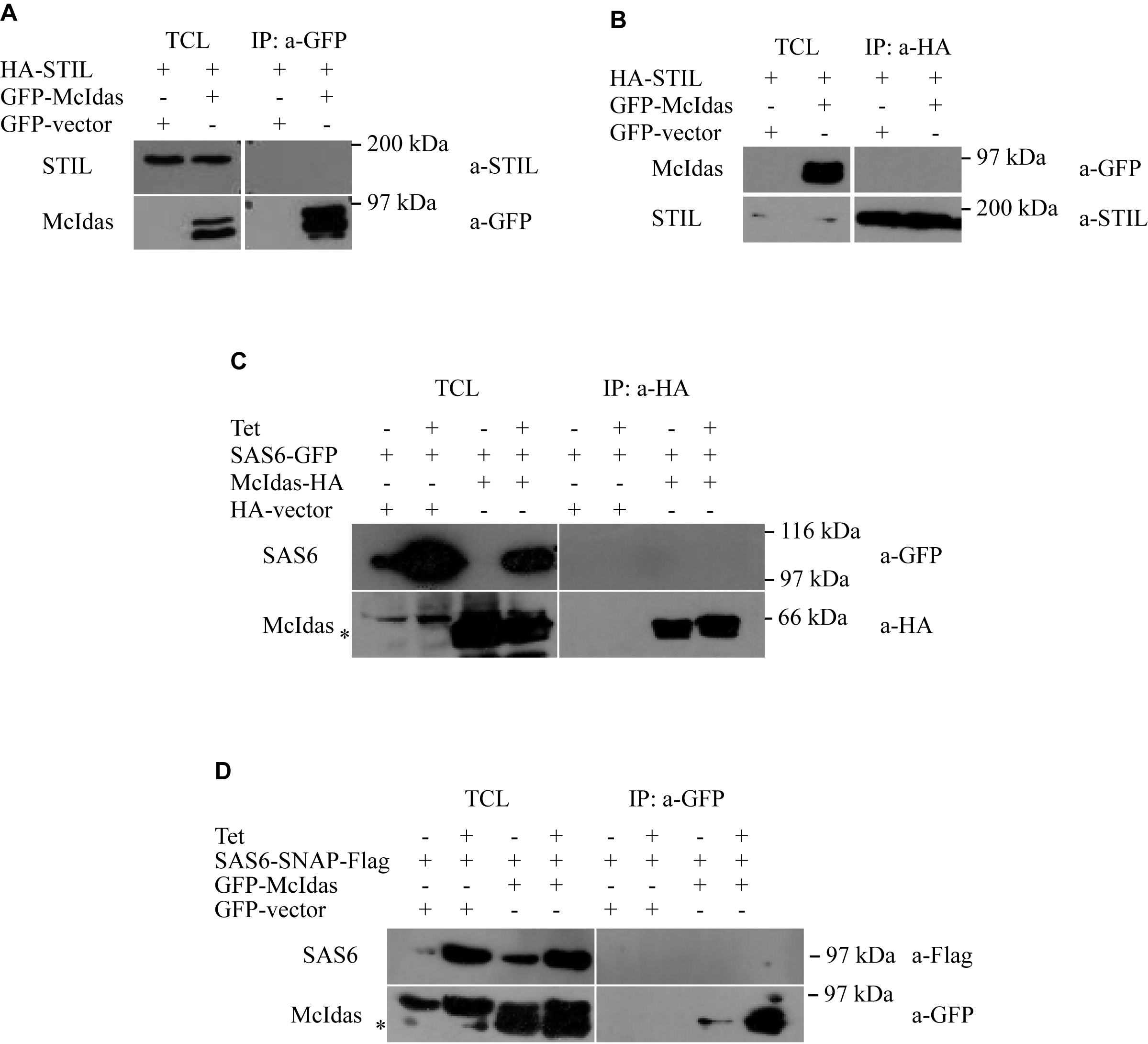

### Supplementary Figure 4

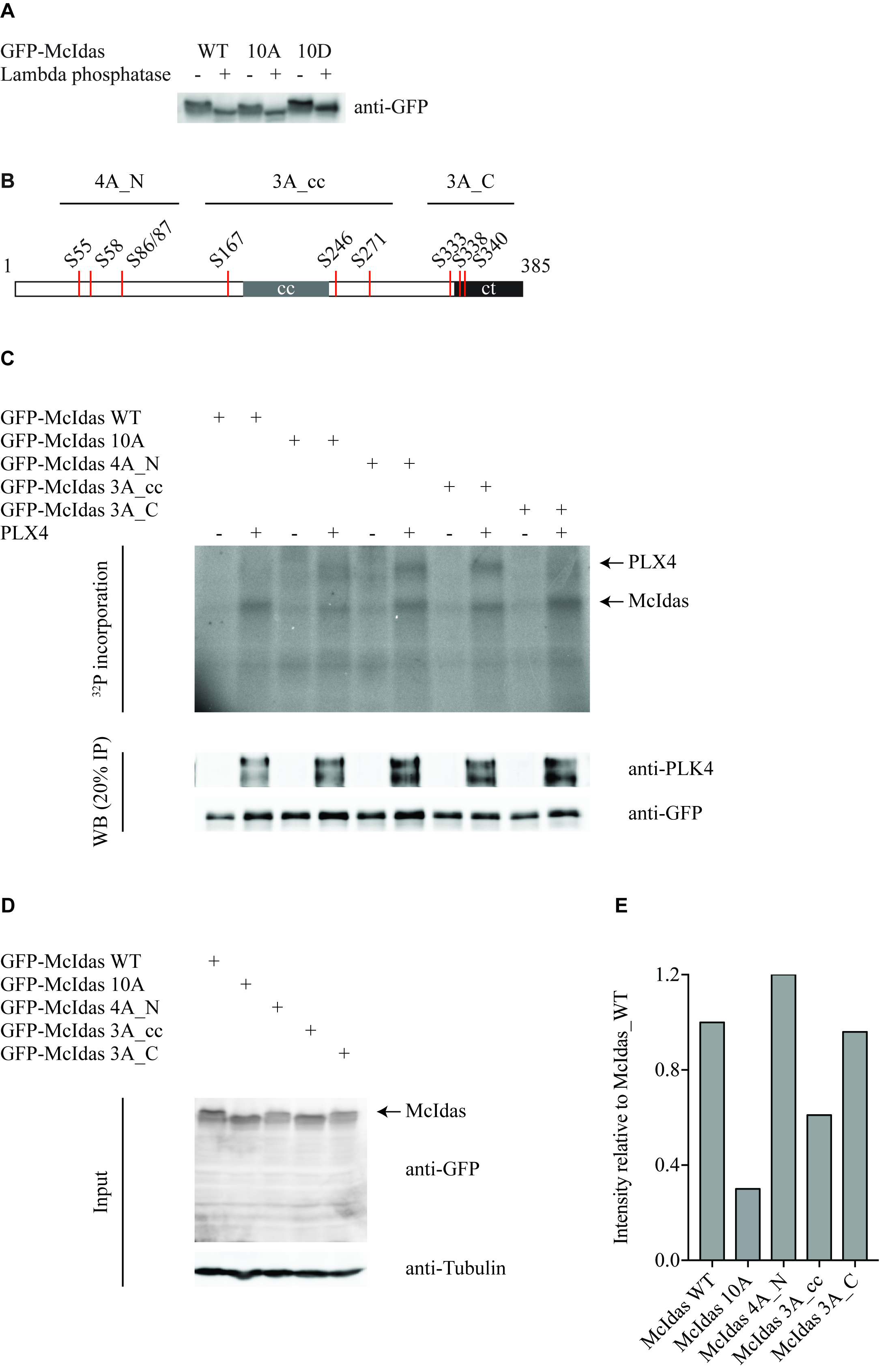
